# Interplay between Anakonda, Gliotactin and M6 for tricellular junction assembly and anchoring of septate junctions in *Drosophila* epithelium

**DOI:** 10.1101/2020.04.27.063131

**Authors:** Thomas Esmangart de Bournonville, Roland Le Borgne

## Abstract

In epithelia, Tricellular junctions (TCJs) serve as pivotal sites for barrier function and integration of both biochemical and mechanical signals. While essential for tissue homeostasis, TCJ assembly, composition and links to adjacent bicellular junctions (BCJs) remain poorly understood. Here we have characterized the assembly of TCJs within the plane of adherens junctions (tAJ) and the plane of septate junctions (tSJ) in *Drosophila* and report that their formation is spatiotemporally decoupled. The assembly and stabilization of previously described tSJ components Anakonda (Aka) and Gliotactin (Gli) as well as the newly reported tSJ proteolipid protein M6, is shown to be a complex process. Aka and M6, whose localization is interdependent, act upstream to locate Gli. In turn, Gli stabilizes Aka at tSJ. Those results unravel a previous unknown role of M6 at tSJ and a tight interplay between tSJ components to assemble and maintain tSJs. In addition, tSJ components are not only essential at vertex as we found that loss of tSJ integrity also induces micron-length bicellular SJs deformations that are free of tensile forces. This phenotype is associated with the disappearance of SJ components at tricellular contacts, indicating that bSJ are no longer connected to tSJs. Reciprocally, SJ components are in turn required to restrict the localization of Aka and Gli at vertex. We propose that tSJs function as pillars to anchor bSJs to ensure the maintenance of tissue integrity in *Drosophila* proliferative epithelia.

## Introduction

Epithelia are tissues fulfilling functions including secretion, absorption, protection, trans-cellular transport and sensing. In addition to these functions, a common feature of epithelia is that they act as mechanical and paracellular diffusion barriers thanks to intercellular junctions. Adherens Junctions (AJs) ensure the mechanical integrity functions [1, 2], while tight junctions in vertebrates (TJ; [3, 4]) and septate junctions in arthropods (SJs; [5–7]), ensure the para-cellular diffusion barrier function. Throughout embryogenesis and then life, epithelial tissues are continuously growing, regenerating or undergoing morphogenesis largely due to cell division. We have contributed to show how cell-cell junctions are remodeled and how tissue integrity is preserved as the epithelial cell undergo cytokinesis. Both the dividing cell and the neighbors are subjected to profound cell shape changes leading to the formation of a bicellular junctions (BCJs) connecting the two daughter cells. At both edges of newly formed BCJ, at the contact with the neighbors, two new tricellular junctions (TCJs) are assembled. Importantly maintenance of mechanical and permeability barrier has to be ensured at the BCJ/TCJ boundary despite their distinct architecture and protein composition. But how this is achieved remains largely unknown.

In *Drosophila*, bicellular SJs forms strands parallel to the plane of the epithelia. SJs are complex assemblies made of multiple protein-protein interactions that are stage and tissue dependent. More than 20 proteins, mostly transmembrane proteins but also GPI-anchored proteins and cytosolic proteins are assembled together to form the SJ [5, 8, 9]. Contactin, Neuroglian (Nrg) and Neurexin-IV (Nrx-IV) are core component proteins of the SJ [10–12]. Nrx-IV intracellular domain binds directly to the FERM domain (Protein 4.1, ezrin, radixin, moesin) of cytoplasmic SJ scaffold protein Coracle (Cora)[13]. Cora forms with Nrx-IV and Nrg a bigger complex with the ion transporter Na^+^/K^+^ ATPase composed of the ATP-α and Nervana 2 (Nrv2) subunits [10, 14]. All of these proteins are interdependent on each other for their proper localization. Proteins of the MAGUK family (membrane-associated guanylate kinase) such as Disc-Large (Dlg) required for cell polarity establishment, colocalize with core SJ components [15]. Fluorescence recovery after photobleaching (FRAP) experiments also revealed that Dlg exhibits a fast recovery (second timescale) whereas the core SJ components Nrx-IV, ATP-α, Nrv2, Nrg and Cora display are recovered with slow kinetics (hour time scale) in matured SJs [16, 17]. The rather stable SJ are running all along bicellular junctions until they reach three cell corners. There, pioneer EM studies in invertebrates revealed that SJ strands make a 90-degree turn when BCJ are abutting TCJ [18–20].

*Drosophila* tricellular junction (TCJ) are composed of the tricellular adherens junction (tAJ) and tricellular septate junction (tSJ). Sidekick (Sdk), a transmembrane protein containing immunoglobulin (Ig) and Fibronectin-type III domains localizing at tAJ, is involved in cell-cell contact rearrangement during development [21–23]. Basal to tAJ, tSJ provide permeability barrier function [24, 25]. tSJs are also key to detect and integrate biochemical and mechanical signals essential for epithelia homeostasis [26, 27]. tSJ is made of Anakonda (Aka), a large transmembrane protein with a tripartite extracellular domain [25, 28], Gliotactin (Gli), a cholinesterase-like transmembrane protein [24, 29], and M6 a glycoprotein of the myelin proteolipid protein (PLP) family [30, 31]. Aka is known as an upstream regulator of tSJ assembly. The tripartite extracellular domain of Aka is proposed to be involved in its stabilization at the vertex [25]. Aka is required for Gli localization at TCJ to ensure tSJ integrity.

Despite recent advances, a global view of TCJ assembly and dynamics is missing. How M6 localize at tSJ and what is its function there remain largely unknown. Whether tSJ and tAJ positioning, assembly and dynamics are coordinated is currently unknown. Also poorly understood are the links between BCJs and TCJs. Knock-down of Aka or Gli was recently reported to cause the basal spreading of Cora, Nrv-2 or Dlg [32]. On the contrary, the GUK domain of Dlg seems to be required for proper localization of Aka and Gli at vertices [32]. While this study suggests a dialog between tSJ and bicellular SJ, the interplay between TCJ and BCJ remains largely underexplored.

In this study, we used time-lapse imaging in the pupal notum of *Drosophila melanogaster* to investigate how tAJ and tSJ are remodeled and formed de novo during cytokinesis. We next examined the hierarchy and interdependency in TCJ component assembly and explored the mechanisms by which tSJ act as pillars. In fact, tSJ is required for SJ integrity maintenance during cytokinesis, regulating SJ morphology as well as SJ core component localization during interphase. In return, we analyzed how SJ proteins regulate the localization of tSJ components at vertices. Our study not only shed new light on TCJ assembly but also reveals the interdependency in TCJ and BCJ components localization, assembly and functions.

## Results

### TCJ *de novo* assembly relies on spatio-temporal polarized mechanism

The notum of the *Drosophila* melanogaster is a proliferative mono-layered epithelium, allowing us to study cell division in time and space. During cytokinesis, one new BCJ interface is established as well as two new TCJs (Figures 1A-1C). To decipher the assembly of TCJ *de novo* (tAJ and tSJ), we imaged the AJ and the midbody using the non-muscle Myosin II light chain tagged with red fluorescent protein (Spaghetti Squash, Sqh∷RFP) together with the tAJ marker Sdk∷GFP or tSJ markers M6∷GFP and Aka∷GFP using time-lapse confocal microscopy. Following actomyosin ring constriction, Sqh is recruited at the new cell-cell interface and in neighboring cells at AJ level (Figure 1A) and finger-like protrusions (FLP) form below at SJ level pointing to the midbody (Figure 1C) as described in [17, 33]. At t=10min, initiation of the formation of new adhesive contacts between daughter cells occurs (Figure 1B) [34–36]. The first signal of Sdk∷GFP appears at the new vertex formed at t=10min 30 sec ± 2 min after anaphase onset (Figures 1B and 1F), concomitantly with the establishment of the new adhesive contacts between daughter cells [34–36].

**Figure 1:**
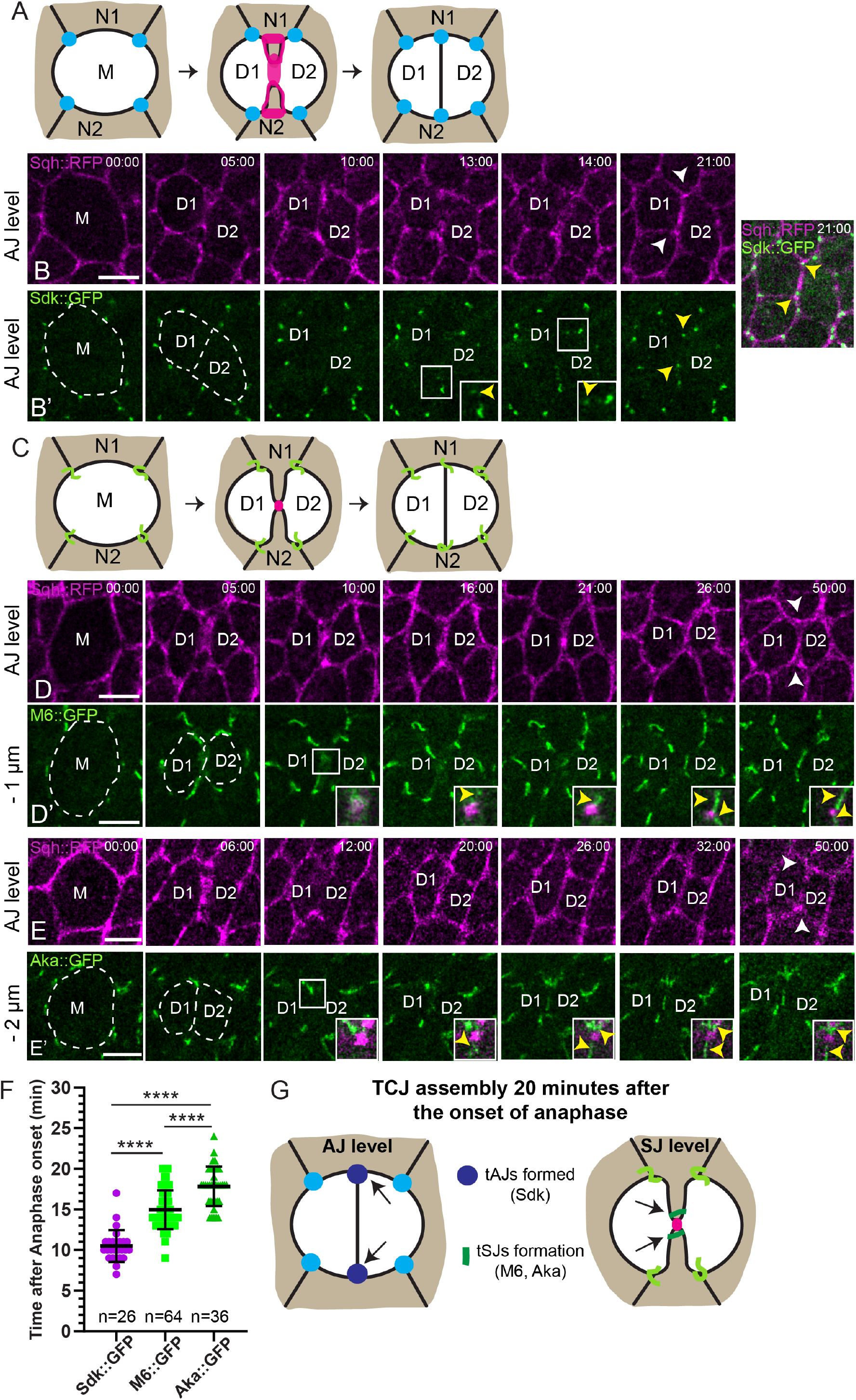
Spatiotemporal analysis of Tricellular junctions assembly during epithelial cytokinesis. (A) Schematic of tAJ assembly during cytokinesis. M represents the mother cell, N the neighboring cells and D the daughter cells. Magenta signal represents Sqh∷RFP enrichment at the new interface and the enrichment on neighboring cells. Blue dots show tAJ. (B, D and E) Time-lapse imaging of Sqh∷RFP^crispr^ (magenta, AJ), Sdk∷GFP (B’; green, tAJ), M6∷GFP (D’; green, tSJ) and Aka∷GFP (E’; green, tSJ) in dividing cells. The mother cell is represented by the M and the daughters by D1 and D2. The white dashed line highlights the divided cell and the two new cells. Yellow arrowheads show the first signal of proteins tagged GFP appearance. White arrowheads show the two new TCJs formed. White squares show high magnifications of Sdk∷GFP (B’) arrival at new vertex or M6∷GFP (D’) /Aka∷GFP (E’) first appearance close to the midbody (magenta). (C) Schematic of tSJ assembly during cytokinesis. M represents the mother cell, N the neighboring cells and D the daughter cells. Magenta signal represents the midbody between the FLP forming at SJ level. Green lines show tSJ. (F) Plot of the mean times of the first appearance after anaphase onset of Sidekick∷GFP (mean = 10 min 30s; purple dots, n = 26 divisions, 4 pupae), M6∷GFP (mean = 15 min; green squares, n = 64 divisions, > 5 pupae) and Anakonda∷GFP (mean = 17 min 50s; dark green triangles, n = 36 divisions, > 5 pupae). Bars show Mean ± SD, **** p < 0.0001, unpaired t test. (G) Schematic of tAJs and tSJs assembly 20 minutes after anaphase onset. Magenta signal represents the midbody between the FLP forming at SJ level. Dark blue dots show new tAJs characterized by Sdk∷GFP and dark green lines show the formation of new tSJs characterized by M6∷GFP and Aka∷GFP, below tAJs. Time is min:sec (B, D and E) with t = 0 corresponding to the anaphase onset. Distances correspond to the position relative to the plane of AJ labeled with Sqh∷RFP^crispr^. The scale bars represent 5μm.

M6 [30, 31] localizes at tSJ at steady state in pupal notum. M6∷GFP first appears in the form of a dispersed signal around 4-5 min after anaphase onset between the two daughter cells at and below midbody level (Figure S1). Then, the first punctate signals appear close to the midbody at t=15min ± 2min 30s after anaphase onset and continue to spread until they form a continuous tSJ strand (Figures 1D and 1F). Moreover, we confirmed that Aka∷GFP punctate signals appear near the midbody at t=17min 50s ± 2min 30s after anaphase onset [17], a step that precedes the formation of a continuous tSJ strands (Figures 1E and 1F).

Thus, assembly of the tAJ and tSJ are spatially and temporally uncoupled (Figure 1G). The presence of tSJ components along the FLP during cytokinesis raises the question of the role of tSJ in FLP maintenance and/or reciprocally the role of FLP on tSJ assembly.

### tSJ components regulate FLP geometry and maintenance at SJ level during cytokinesis

We then used Sqh∷RFP in combination with ATP-α∷GFP in *aka*^*L200*^ cells using clonal mosaic analysis to study the role of Aka in FLP formation and/or maintenance during cytokinesis. In wild-type conditions, FLPs form as the consequence of actomyosin cytokinetic ring contraction, 1μm below AJ (t = 5min; Figures 2A-2B). 10 minutes after anaphase onset, FLPs were pointing to the midbody and displayed a characteristic “U” shape (Figures 2A’-2B). Midbody was centered in 71% of the cases resulting in the formation of two FLPs of similar length (Figures 2A’, 2B and 2E) that persist more than 20 minutes after anaphase onset (Figures 2A-2B). Upon loss of Aka, actomyosin cytokinetic ring contraction also led to the formation of FLP (t=5 min; Figures 2C-2D). Despite the initial symmetry of the ring constriction, the length of the two FLPs was unequal (62% of cases compare to only 29% for wild-type cells; Figures 2C-2E) and midbody is off-centered. In addition, the FLP exhibits a narrower width compared to control, and their tip formed a loop shape (Figures 2C’, 2C’’ and 2D). Moreover, the midbody is found more basal upon loss of Aka (between 1 and 2μm below AJ, Figures 2C-2C’’’) in accordance with the faster basal displacement as described previously [17]. Finally, we observed that ATP-α∷GFP signal is inhomogeneous in those FLPs compared to wild-type conditions (Figures 2C’-2D). Because one *aka*^*L200*^ cell adjacent to a vertex leads to tSJ disruption [25], we looked at a wild-type cell dividing between one wild-type cell and one *aka*^*L200*^ cell (Figures S2A-S2B). Midbody was off-centered in 57% of cases (Figures S2A-S2C). These results indicate that Aka ensures the symmetry in FLP formation as well as the SJ components distribution, suggesting a role of Aka in regulating BCJ (see below) (Figure 2F).

**Figure 2:**
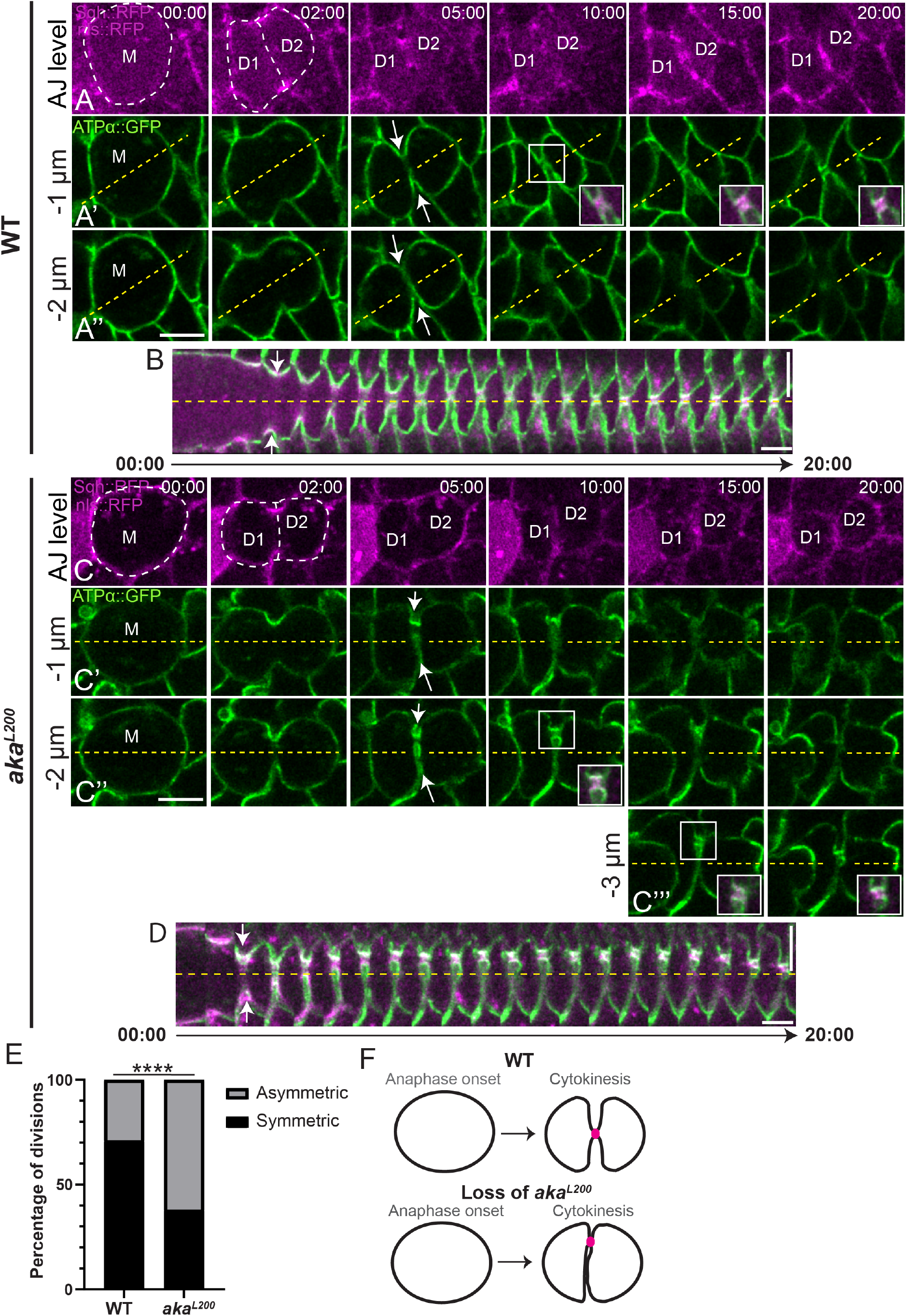
tSJ integrity is required to ensure FLP formation during cytokinesis. (A-D) Time-lapse imaging of FLP using Sqh∷RFP^crispr^ (magenta, AJ) and ATPα∷GFP (green, SJ) in wild type (marked by nls∷RFP; A-B) and *aka*^*L200*^ (loss of nls∷RFP; B-D) dividing cells, from plane view (A-A’’ and C-C’’’) and with a kymograph representation (B and D). The mother cell is represented by the M and the daughters by D1 and D2. The white dashed lines highlight the divided cell and the two new cells. White arrows indicate FLP formation at SJ level. Yellow dashed lines define symmetry axis of the cell. White squares show high magnification of midbody (magenta) linking protrusion like fingers. (E) Histogram representing the number of symmetric (Black) and asymmetric (Grey) FLP formation during cytokinesis in wild type (symmetric = 71% ; asymmetric = 29% ; n = 14 divisions, 3 pupae) and *aka*^*L200*^ cells (symmetric = 38% ; asymmetric = 62% ; n = 21 divisions, > 5 pupae). **** p < 0.0001, Fisher’s exact test. Schematic representation of the FLP formation upon cytokinesis. Loss of tSJ integrity leads to abnormal FLP shape and asymmetrical midbody formation at the new cell-cell interface. The horizontal scale bars represent 5μm (A and C), the vertical scale bars represent 5μm (B and D) and the horizontal scale bars represent 1min (B and D).

In contrast to SJ, no defects were seen at vertex at AJ level upon loss of Aka throughout cytokinesis. Together with the fact that the recruitment of Sdk and Aka/M6 are spatially and temporally decoupled during cytokinesis (Figure 1), these data suggest that tSJ and tAJ assembly and perhaps functions are uncoupled.

### Interplay between tAJ and tSJ components

We first analyzed the consequence of loss of Sdk on tSJ. We used the homozygous viable null allele *sdk*^*Δ15*^. We observed no differences of Aka signals between wild-type and *sdk*^*Δ15*^ pupae (Figures 3A-3B’). Conversely, we studied the loss of tSJ integrity using the *aka*^*L200*^ mutant on Sdk localization. No differences in the localization of Sdk were observed between wild-type and *aka*^*L200*^ cells (Figures 3C-3C’). These data indicate that the localization, recruitment and/or stabilization of Sdk and Aka at TCJ are independent one of each other (Figure 3G).

**Figure 3:**
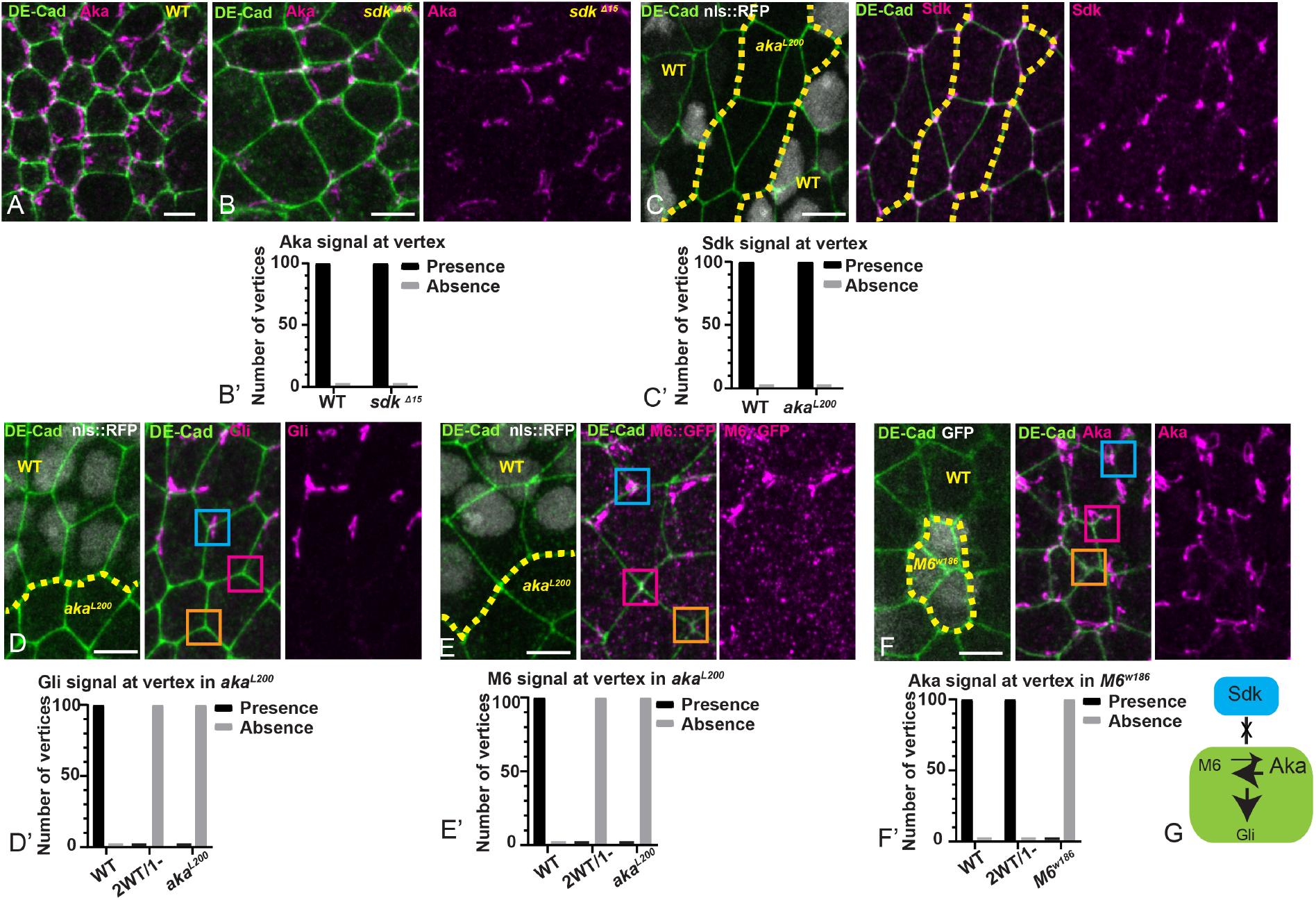
Aka and M6 are required for tSJs and dispensable for tAJs integrity. (A-B) Localization of Aka (anti-Aka, magenta) in wild type and *sdk*^*Δ15*^ pupa. (B’) Histogram representing the percentage of presence (black) or absence (gray) of Aka at the vertex in wild type (presence n = 100, absence n = 0) and *sdk*^*Δ15*^ (presence n = 100, absence n = 0) (n = 100 vertices, 3 pupae) (C-F) show nota between 16h-18h APF stained for DE-cad (anti-DE-cad, green) after heat-shock to induce clone of wild type (C, D, E nls∷RFP positive; F, GFP negative) and mutant cells (C, D, E nls∷RFP negative; F, GFP positive) for TCJ components. (C) Localization of Sdk (anti-Sdk, magenta). The dashed yellow line separates wild type and *aka*^*L200*^ cells. (C’) Histogram representing the percentage of presence (black) or absence (gray) of Sdk at the vertex in wild type (presence n = 100, absence n = 0) and *aka*^*L200*^ (presence n = 100, absence n = 0) (n = 100 vertices, > 5 pupae). (D) Localization of Gli (anti-Gli, magenta). The dashed yellow line separates wild type and *aka*^*L200*^ (mutant) cells. Gli is enriched at the TCJ at wild-type vertex (blue square) and disappeared at a vertex of three *aka* mutant cells (orange square) but also when Aka is lost in only one of the three cells participating in the vertex (pink square). (E) Localization of M6 (M6∷GFP + anti-GFP, magenta). Wild-type and *aka*^*L200*^ cells are separated by the dashed yellow line. M6 is enriched at the TCJ at wild-type vertex (blue square) and disappeared upon loss of Aka from one cell adjacent to the vertex (pink square) or at a vertex of three *aka*^*L200*^ cells (orange square). (F) Localization of Aka (anti-Aka, magenta). Wild type and *M6^w186^* mutant cells are separated by the dashed yellow line. Aka is enriched at the TCJ at wild-type vertex (blue square) and upon loss of M6 from one cell adjacent to the vertex (pink square) but disappear at vertex of three *M6^w186^* mutant cells (orange square). (B’, C’, D’, E’ and F’) Histograms representing the percentage of presence (black) or absence (gray) of Gli/M6/Aka at the vertex between 3 wild type cells (presence n = 100, absence n = 0), 2 wild type and 1 *aka*^*L200*^ cell (D’ and E’; presence n = 0, absence n = 100) or 1 *M6^w186^* cell (F’; presence n = 100, absence n = 0) or 3 *aka*^*L200*^ cells (D’ and E’; presence n = 0, absence n = 100) or 3 *M6 ^w186^* cells (F’; presence n = 0, absence n = 100) (n = 100 vertices, > 5 pupae for each experiment). Schematic of the interplay of TCJ components. The scale bars represent 5μm. All images are maximum projection.

We next investigated the relationship between the three tSJ components. Loss of Aka leads to Gli and M6∷GFP disappearance from tSJ, even when only one cell contributing to the vertex is mutant for *aka* (Figures 3D-3E’), thereby confirming that Aka is an upstream regulator in the notum, as previously described in *Drosophila* embryo [25]. We next investigated the role of M6 on Aka and Gli localization and found that loss of M6 leads to Aka (Figures 3F-3F’) and Gli (Figures S3A and S3B) disappearance at vertex. Therefore, M6 is like Aka an upstream regulator of tSJ assembly. However, whereas loss of Aka from only one cell contributing to the vertex leads to a complete loss of M6 at tSJ, loss of M6 from one cell contributing to the vertex triggers the reduction in Aka signal (Figures 3F-3F’) or Gli signal (Figures S3A and S3B) rather than a complete disappearance.

### Gli stabilizes Aka at tSJ

Because Gli activity is dispensable for Aka localization at tSJ in embryo [25], we tested whether this was also the case in the pupal notum. Unexpectedly, we found a decrease in the intensity and the surface occupied by Aka signal in *Gli*^*dv3*^ mutant cells (Figures 4A-4B’’). We next tested the possibility that the stabilization of Aka at tSJ is modified upon loss of Gli using FRAP. Our experiments revealed that upon loss of Gli, the rate of fluorescence recovery of Aka∷GFP is higher (t_1/2_ = 318s) compared to wild-type conditions (t_1/2_ = 361s) (Figures 4C-4D). In addition, immobile fraction of Aka∷GFP drastically dropped to 0.30 compared to 0.75 in wild-type conditions, 20 minutes after photobleaching (Figure 4D).

**Figure 4:**
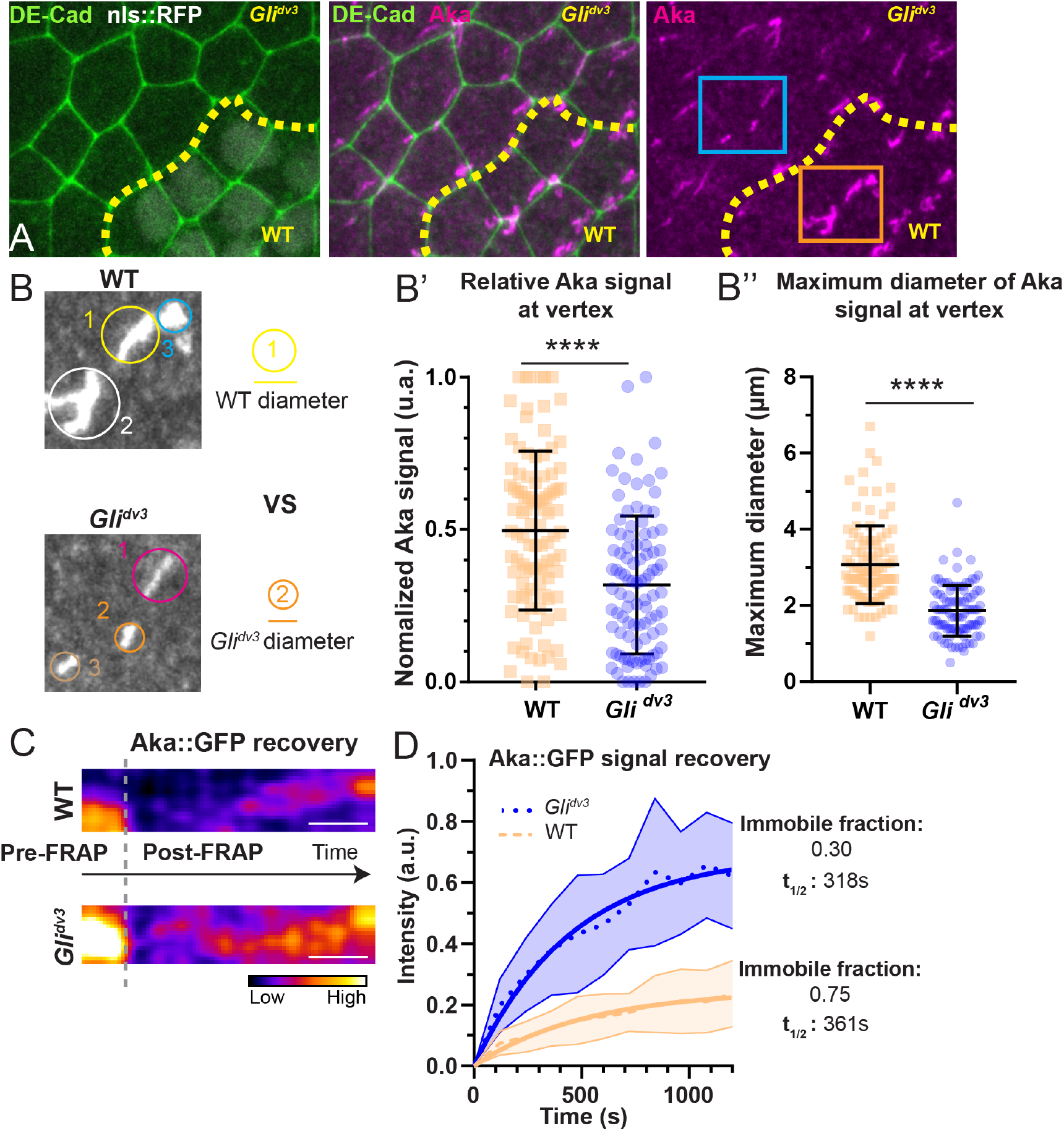
Gli stabilizes Aka at the tSJ. (A) Localization of Aka (anti-Aka, magenta) in notum marked by DE-cad (anti-DE-cad). Wild-type (nls∷RFP positive) and *Gli*^*dv3*^ cells (nls:RFP negative) are separated by the dashed yellow line. Orange and blue squares show magnification area for B. (B) Magnification of panel A pictures depict Aka signal in wild-type (orange square) and *Gli* cells (blue square). The diameter of the circle including the total Aka signal is measured, extracted and compared between wild type and *Gli*^*dv3*^ cells. (B’) Plot of the of normalized Aka signal at vertex in wild type (orange squares) and *Gli*^*dv3*^ cells (blue circles). (n = 100 vertices each, > 5 pupae) (B’’) Plot of the diameters (μm) of circles including maximal Aka signal at vertex in wild type (orange squares) and *Gli*^*dv3*^ cells (blue circles). (n = 100 vertices each, > 5 pupae). Bars show Mean ± SD, **** p < 0.0001, unpaired t test. (C) Kymograph of the bleached region for Aka∷GFP in wild type and *Gli*^*dv3*^ cells. A calibration bar shows LUT for gray value range. Scale bars show 5min. (D) Plot of Aka∷GFP fluorescence recovery as a function of time for the conditions described in (C). Wild type: n = 11 FRAP experiments, 4 pupae; *Gli*^*dv3*^: n = 12 FRAP experiments, 5 pupae. Data are mean ± SD. Solid line shows a simple exponential fit.

Therefore, while being dispensable to localize Aka at tSJ, Gli influences the time of residency of Aka, revealing an interplay between Aka, Gli and M6 (Figure 3G).

### tSJ ensures SJ integrity at both bicellular junctions and vertices

Our time-lapse analyses upon loss of Aka also revealed defects in the integrity of bSJ in interphase. Indeed, loss of Aka caused the appearance of 1 to 3 micron-long membrane deformations labelled using GAP43∷mCherry (Figures S4A-S4B). These membrane deformations are located within the plane of SJ, positive for the SJ core components including Cora (Figure 5A) ATP-α and Nrx-IV (Figures 5B and 5C), as well as Dlg (Figure 5D). This phenotype was not only observed using several *aka* mutant alleles (data not shown), upon silencing of Aka (Figure S4C) but also upon loss of Gli (Figure S4D) indicating that loss of Aka or its stability at tSJ is associated with SJ membrane deformations. Using Nrx-IV as a marker, we observed that many deformations are found apical, close to and/or within the plane of DE-cad albeit not colocalizing with it (Figures 5E-5H’). We also observed Nrx-IV signal disappearance or drastic decrease from vertices in *aka*^*L200*^ cells (Figures 5I and 5I’). While similar results were observed for ATP-α∷GFP (Figure S4E), in contrast Dlg remained present at the vertex (Figures 5I and 5I’). This data argue for a depletion of SJ core components from the vertex. The loss of Gli induced similar results on Nrx-IV presence at vertex (Figures S4F and S4G). These results raise the question of how and why loss of tSJ integrity prevents the presence of SJ core components at the vertex (Figure 5J) and cause SJ deformations.

**Figure 5:**
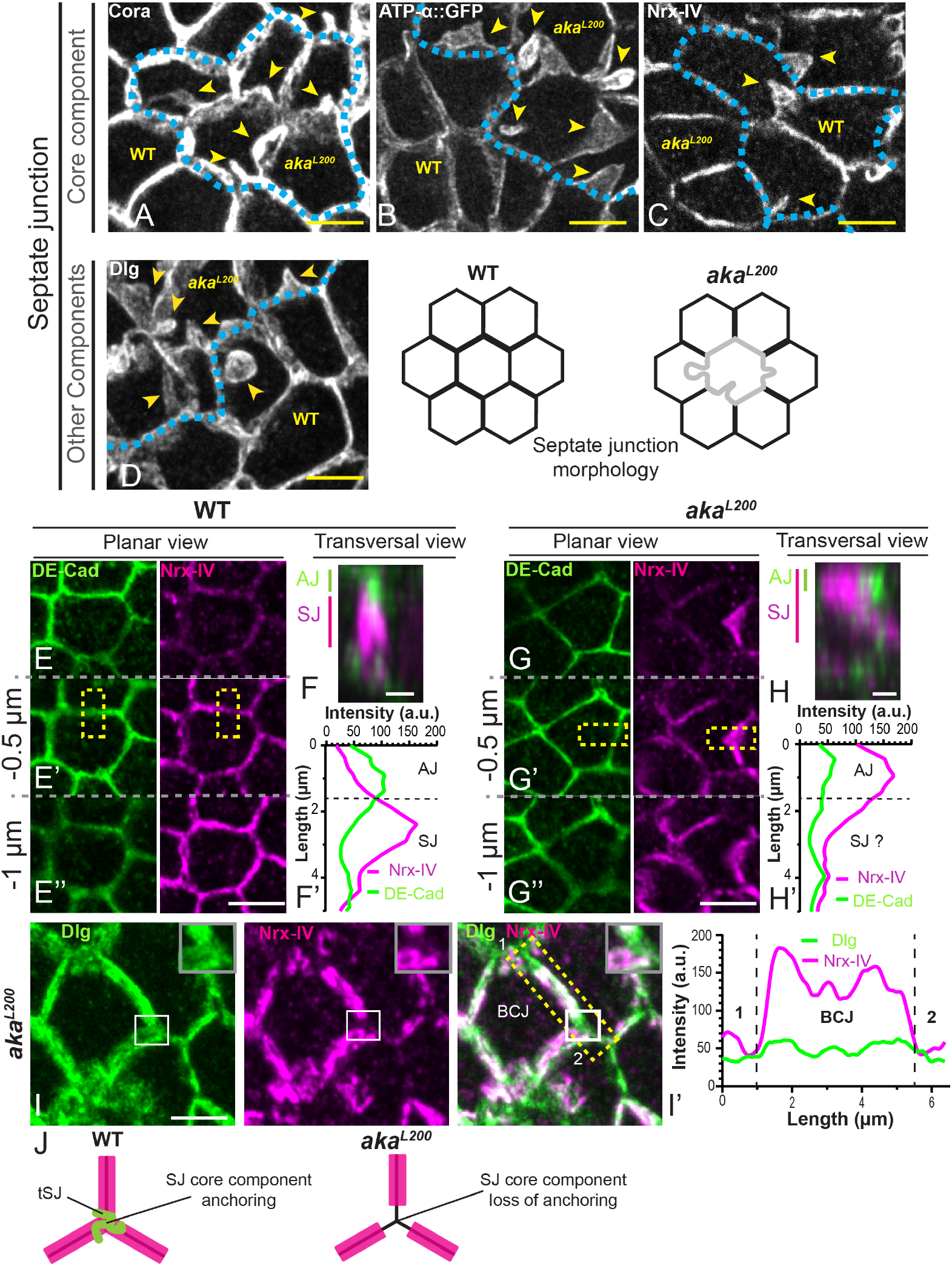
tSJs regulate the shape of SJ and ensure the anchoring of SJ core components at the vertex. (A-D) Analyses of SJ morphology via core components Cora (anti-Cora), ATP-α (ATP∷GFP), Nrx-IV (anti-Nrx-IV) and other SJ component Dlg (anti-Dlg) in wild type and *aka*^*L200*^ cells. Yellow arrowheads indicate micrometric size deformations of SJ. The dashed blue line separates wild type and *aka*^*L200*^ cells. (E-H’) Localization of Nrx-IV (anti-Nrx-IV, magenta) in notum marked by DE-cad (anti-DE-cad, green). (E-E’’’ and G-G’’’) show different planar sections separated by 0.5μm in wild type and *aka*^*L200*^ cells. (F and H) show maximum projection of the transversal section depicted by the yellow rectangles. (F’ and H’) Plots representing DE-cad (green line) and Nrx-IV (magenta line) signals as a function of cell transversal length represented in transverse sections. Black dashed lines show the presumptive boundary between AJ and SJ. (I) Localization of Dlg (anti-Dlg, green) and Nrx-IV (anti-Nrx-IV, magenta) in *aka*^*L200*^ cells after maximal projection. White squares show high magnification of a vertex in *aka*^*L200*^ cells. Yellow dashed rectangle shows the line scan used between two vertices represented by 1 and 2 to obtain (I’) Plot representing Dlg (green line) and Nrx-IV (magenta line) signals as a function of the length of cell-cell boundary. (J) Scheme showing the loss of *aka*^*L200*^ effect on SJ core component. Magenta lines show SJ core component, green lines the tSJ and black lines the membrane as well as Dlg protein. The scale bars represent 5μm or 1μm in (F and H).

### Once formed, SJ deformations are fixed in time and space

We then used time-lapse microscopy to live image the biogenesis of these SJ-containing membrane deformations during interphase. We observed that the induction of membrane deformation occurs at low velocity (Figures S5A and S5B) and that they remain at the same location for many hours (Figure S5C). In addition, the SJ-containing membrane deformation elongated (Figures S5D and S5E) while the total BCJ length increases marginally (Figure S5D). This data suggests that extra amount of membrane and new SJ components are brought at the level of the deformation.

Since SJ-containing membrane deformations localize in part at AJ level and that AJs are the sites of acto-myosin driven forces of the tissue [1, 2], we thought to probe if mechanical forces were involved in the stabilization of these deformations. Using two-photon laser-based nano ablation, we cut the tips of deformations to reveal possible pulling or pushing forces (Figure 6A). No recoil or change in the shape of deformation were observed upon ablation (Figure 6A). Then, we probed lateral forces by cutting the bSJ close the deformation and again, revealed no deformation shape changes after ablation (Figure 6B), even by mechanically isolating the mutant cell with a circular cutting (Figure S5F). In accordance with those results, nor Sqh or Actin marked by phalloidin were enriched in those deformations, excluding the possibility that AJ components (DE-cad) or actomyosin stabilize them (Figures S5G and S5H). The only time the membrane deformation was remodeled was during mitosis, when the cell has rounded up (Figure S5I). Nonetheless, during cytokinesis, the former SJ-containing membrane deformation reappears at the location it was before cell division (Figure S5I). Then, we tested if changes in SJ components dynamics could be responsible for the induction of those deformations, by FRAP approach on ATP-α∷GFP. Wild-type cells exhibited slow turnover at bSJ (t_1/2_ = 280s) with an immobile fraction of 0.67 18 min after photobleaching (Figures 6C and 6F). In *aka*^*L200*^ cells, the kinetics of recovery were similar to wild-type condition (t_1/2_ = 294s) but exhibits a slightly increase of the immobile fraction of 0.76 18 min after photobleaching (Figures 6D and 6F). Moreover, FRAP on SJ containing membrane deformations revealed a similar t_1/2_ = 296s but a higher immobile fraction of 0.82 18 min after photobleaching (Figures 6E and 6G). Therefore, loss of tSJ leads to an increased residency time of ATP-α∷GFP. This might explain why the steady state levels of ATPα∷GFP signal is higher in membrane deformations.

**Figure 6:**
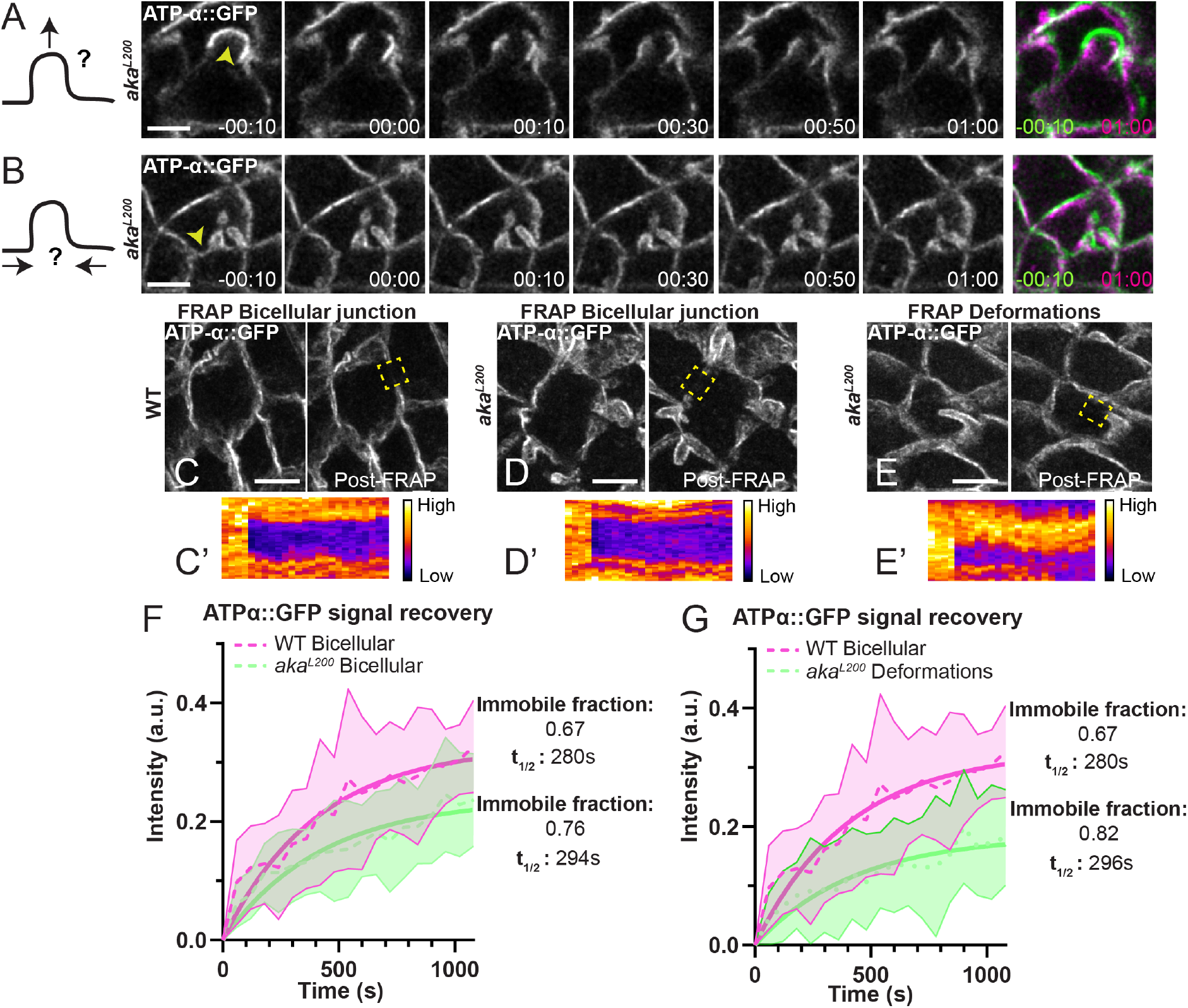
SJ deformations stabilization does not rely on mechanicals forces and SJ proteins exhibits increased stability within SJ deformations. (A and B) Bi-photonic laser-based nanoablation of SJ deformations in *aka*^*L200*^ cells expressing ATP-α∷GFP (white). Green and magenta pictures represent SJ deformations 10s before and 1min after ablation respectively. Yellow arrowheads show the ablation area. (C-E) Example of FRAP experiment of ATP-α∷GFP at wild-type bicellular junction, *aka*^*L200*^ bicellular junction or SJ deformations in *aka*^*L200*^ cells and their associated kymographs (C’-E’). Yellow dashed rectangles show FRAP area. (F and G) Plot of ATP-α∷GFP fluorescence recovery as a function of time for the conditions described in (C-E). Wild type bicellular junction: n= 8 FRAP experiments, 4 pupae; *aka*^*L200*^ bicellular junction: n= 6 FRAP experiments, 4 pupae. *aka*^*L200*^ SJ deformations: n= 9 FRAP experiments, 4 pupae. Data are mean ± SD. Solid line shows a simple exponential fit. The scale bars represent 5μm.

### bSJs restrict the localization of tSJ components at vertices

Loss of tSJ leads to detachment of bSJ from vertex and changes in the dynamics of recovery of SJ core components in membrane deformations, suggesting a possible physical connection between tSJ and bSJ. A prediction of this hypothesis is that loss of bSJ core components would affect tSJ components localization. This prompted us to study the consequence of loss of function of bSJ on the localization of tSJ. We found that depletion of the SJ core component Cora leads to the spreading of Aka along bSJ (Figures 7A-7C’). The lateral spreading of Aka was not due to an increase in the length of bicellular junction adjacent to the vertex (Figures 7D-7E). Similar results were obtained upon depletion of Nrx-IV (Figures S6A-S6C’). Finally, the spreading of Gli was also observed in Nrx-IV RNAi context (Figures S6D-S6F’) and Cora RNA-i (data not shown). Together, our results demonstrate that bSJ integrity is required to confine tSJ components localization at the vertex.

**Figure 7:**
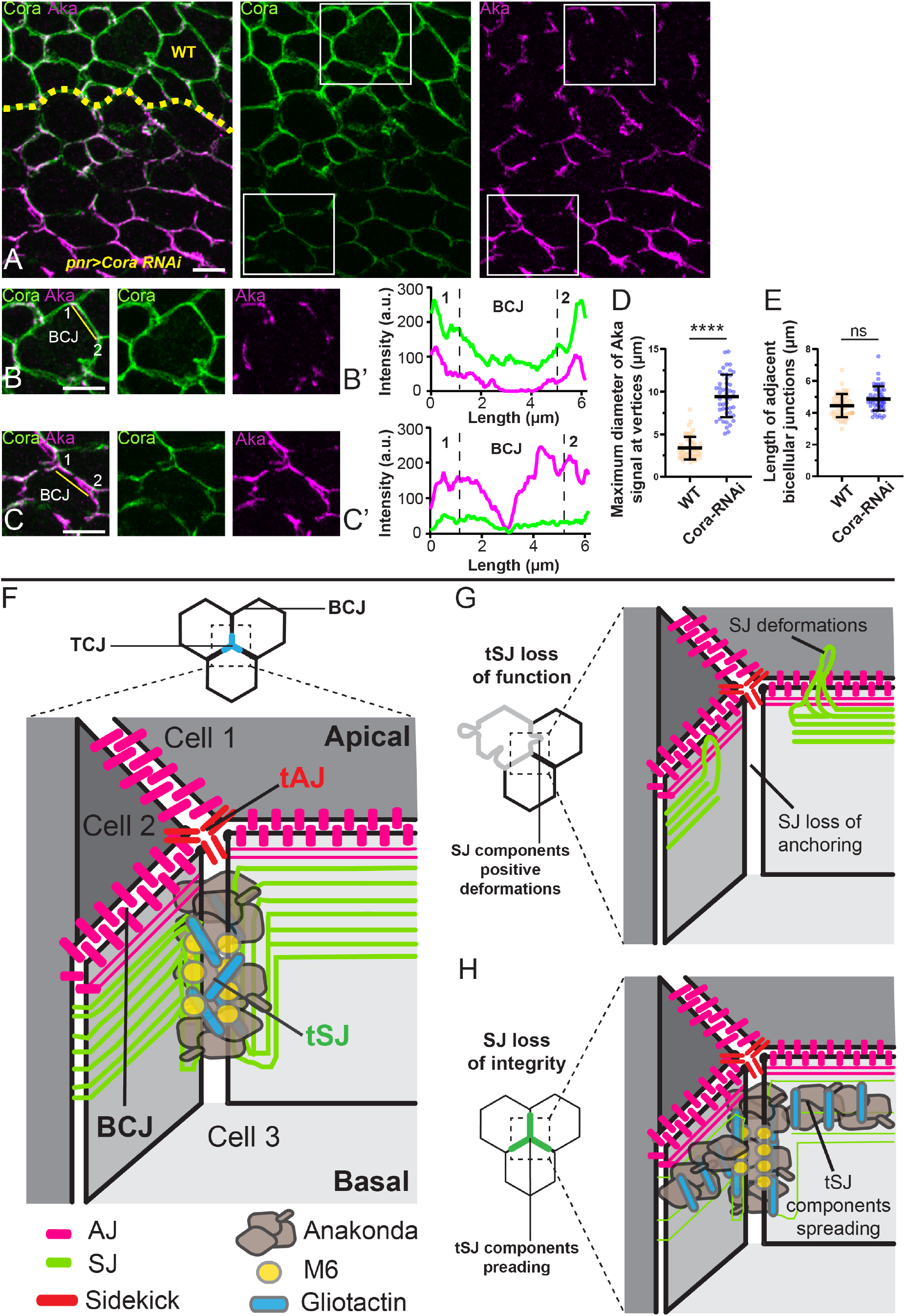
SJ integrity is required to confine tSJ components at vertex. (A) Localization of Aka (anti-Aka, magenta) in cells marked by Cora (anti-Cora, green) in wild type and cells expressing UAS∷Cora-RNAi under pnr∷Gal4 control. The dashed yellow line separates wild type and Cora-RNAi cells. Aka spreads at the BCJ upon knock-down of Cora. White squares show (B and C) magnification of wild type and knock-down cells for Cora. Yellow lines show the line scan used to obtain (B’ and C’) Plots representing Cora (green line) and Nrx-IV (magenta line) signals as a function of the length of cell-cell boundary in wild type and Cora-RNAi cells respectively. (D) Plot of the diameters (μm) of circles including maximal Aka signal at vertex in wild type (orange squares) and Cora-RNAi cells (blue circles). n= 50 vertices, > 5 pupae. (E) Plot of the length (μm) of BCJ adjacent of wild type vertices (orange squares) and Cora-RNAi vertices (blue circles). n= 50 BCJ, > 5 pupae. Bar show Mean ± SD, **** p < 0.0001, ns: non-significant, unpaired t test. (F) Schematic of the TCJ general organization in *Drosophila* notum. (G) Upon loss of tSJ integrity, SJ strands are no longer anchored on tSJ complex, leading to loss of SJ components presence at vertex and SJ deformations inductions in both AJ and SJ level. (H) Upon silencing of SJ core components, Aka and Gli spread at BCJ and are no longer restricted at vertex. The scale bars represent 5μm.

## Discussion

In this study, we first described how TCJ are assembled during cytokinesis and provided evidence that the recruitment, localization and stabilization of tAJ and tSJ are uncoupled and not interdependent. Among tSJ, Aka and M6 were found to be upstream regulators of tSJ while Gli stabilized Aka at tSJs. Moreover, we uncover that tSJ components are essential to maintain SJ homeostasis both in interphase and during mitosis. During cytokinesis, Aka regulates the geometry and length of FLP as well as the homogeneity in SJ proteins distribution. In interphase cells, tSJs control the morphology of bSJs and the presence of core SJ components at vertex. Conversely, SJ integrity is required for tSJ components to be confined at vertices. Based on these results, we propose a mutual dependency model in which tSJ acts as a pillar to anchor SJ strands at vertices while SJ core components acting to restrict tSJ components at vertices (Figures 7F-7H).

### TCJ de novo assembly and components interplay

Our study of TCJ assembly showed that like AJ and SJ, tAJ is established first, prior to tSJ. *De novo* formation of AJ occurs concomitantly to Sdk recruitment at vertices (this study; [34–36]), suggesting a coupled mechanism to ensure AJ mechanical integrity at both bicellular and tricellular junction. In contrast, as loss of Sdk did not prevent Aka localization and reciprocally (this study, [22]), assembly of tAJ and tSJ appears to be spatially and temporally uncoupled.

Once tAJ are formed, tSJ components begin to be recruited at FLP level, with M6 detected before Aka. It is interesting to note that, during cytokinesis in vertebrates, Tricellulin and Lipolysis-stimulated lipoprotein receptor LSR, the main components of tight Tricellular junction (tTJ) [37–39], are also recruited with different timing [40], suggesting similarities for tSJ and tTJ assembly. While, the significance of the differences in the kinetics of recruitment of M6 and Aka is yet unknown, it is conceivable that they are recruited at the same time but that their amounts, possibly reflecting their stoichiometry, differ, leading to an earlier detection of M6 compared to Aka. Another possibility could be that M6 is recruited first to initiate Aka and Gli stability during tSJ establishment.

Our study also investigated the hierarchy in tSJ assembly. We first confirmed that Aka is, as described previously in *Drosophila* embryo [25], an upstream regulator of tSJ assembly, required for localizing Gli. M6 is a tumor suppressor gene [30] which was reported to localize at tSJ. Here we show that Aka and M6 are interdependent for their localization at tSJ, and are acting upstream of Gli in the tSJ assembly pathway. Nonetheless, our analyses of clone borders suggest some differences in the requirement of Aka and M6 in tSJ assembly. Loss of Aka from one cell adjacent to the vertex leads to loss of M6 and Gli but loss of M6 from only one cell of the vertex does not promote the complete Aka and Gli disappearance at vertex. Structure function analyses and mode of subcellular localization of M6, a small four-pass transmembrane proteolipid protein will help understanding its activity, stoichiometry, and relationship with Aka (see companion paper by Wittek et al). Also, it will be interesting to determine whether the mammalian orthologous of M6, enriched in CNS [41, 42] also localizes at TCJ and exerts similar function in vertebrates.

In addition to unravel the function of M6 in tSJ assembly, our study revealed that Gli is required for the maintenance of Aka localization at tSJ. Thus, Gli is not simply a downstream effector of Aka and M6, but it plays an active role in recruiting or stabilizing Aka, and likely M6 at the vertex, indicating a complex interplay between Aka, M6 and Gli in regulating tSJ assembly, stability and function. Loss of LSR in vertebrates leads to the spreading of Tricellulin, resembling the loss of Aka on Gli [38]. Whether or not Tricellulin plays a role on LSR stability remains to be determined.

### tSJ role on SJ integrity

Transmission electron microscopy (TEM) revealed that vertices in invertebrates are intercellular space spanned vertically with diaphragms [18–20]. SJ were also characterized by TEM as belt-like strands of septa [18–20, 43]. Those strands run parallel to the apical surface at intercellular space and start to turn vertically close to the vertex, forming limiting septa in contact with diaphragms that are parallel now to the vertex axis. In fluorescent microscopy, SJ strands looks like either accordion or well defined strand depending on *Drosophila* developmental stage [16, 44]. tSJs display a snake like shape, making hard to reconcile TEM and fluorescent based model. Moreover, SJ proteins partially colocalize with tSJ proteins (this study; [32]), suggesting that SJs and tSJs are interacting physically each other’s. Our results are compatible with a model in which Gli interacts directly with SJ components proteins. Another hypothesis could state that the loss of Aka stability by loss of Gli triggers SJ core components exclusion from the vertex. If so, the effect of loss of Gli on Aka stability might be indirectly mediated since co-immunoprecipitation experiments showed negative results about Gli and Aka interaction [25]. Gli is part of Neuroligin family [45], a set of proteins involved in synapse formations [46] and synaptic transmission [47, 48]. Moreover, recent studies highlighted physical interaction between MGDA proteins and Neuroligin [49–51]. MGDA proteins are composed of six N-terminal Ig domains and one fibronectin type III domain [52], which looks like Nrg organization, displaying also six N-terminal Ig domains followed by five fibronectin type III domain [53]. MGDA are reported interacting with Neuroligin via their first two Ig domains [49–51]. Based on our results and knowing that Nrg formed a complex with Cora, ATP-α and Nrx-IV [6, 10], the link between tSJ and SJ could be Gli binding Nrg. Therefore, these data reveal the potential link between SJ and tSJ, making tSJ the pillar required to anchor SJ strands.

### tSJ loss of integrity lead to cellular SJ defects compensation

Furthermore, upon loss of tSJ components, large SJ membrane deformations form. They exhibit a slower ATP-α∷GFP recovery following photobleaching, a quasi-fixed position across time, no actomyosin cytoskeleton enrichment, and no detectable mechanical forces being implicated in their stabilization. However, during cytokinesis when actin cytoskeleton is remodeled and cortical tension increases [54, 55], SJ deformations lost their shape and followed the cell curvature prior to anaphase onset. Therefore, SJ deformations can be remodeled under high cortical tension. Surprisingly, we could identify their previous localization since SJ enrichment persisted during mitosis. After anaphase, SJ deformations take back their position and shape, suggesting a “form memory”. Epithelial cells lacking CrebA showed excess of membrane and SJ components [56]. Also, another recent study [44] highlighted that subperineurial glial cells lacking SJ integrity compensate by overexpressing SJ components and making more cellular membrane. By analogy, it is conceivable that the loss of SJ strands anchoring at tSJ leads to cell detection of SJ permeability defect. Then, more SJ proteins and membrane components are targeted to plasma membrane. If so, the slow turnover of SJ proteins could not handle the extra amount of SJ proteins leading to SJ proteins local enrichment, elongation of SJ with the making of deformations, preventing even more their basal spreading or recycling. However, how cell detects SJ permeability defects and what molecular processes are involved in SJ deformations remains elusive at present.

### SJ integrity restrict tSJ proteins at vertex

The mutual exclusion of SJ and tSJ combined with the pillar model proposed above raise the question about origins of tSJ localization. Inducing loss of SJ integrity leads to Aka and Gli spreading along SJ as shown here upon Cora depletion. One hypothesis could be that the space availability made by SJ depletion allows tSJ components to move at this location. Another explanation could be that tSJ components roles are multiple. One is to anchored SJ strands at vertices to ensure their shape and integrity. Another one could be that they also play a role in permeability function, by localizing SJ components in close vicinity of vertex or by playing a filter role themselves. Therefore, their spreading at SJ upon SJ loss of integrity could be a compensatory mechanism.

Based on the conservation of epithelial barrier functions, it is tempting to speculate that the relationship we here uncover between TCJ and BCJ components might also oapply to vertebrates to ensure maintenance of epithelial tissue integrity.

## Acknowledgements

We thank C. Klämbt, S. Luschnig, A. Martin, J. Treisman, A. Uv, T. Xu, the Bloomington Drosophila Stock Center, the Vienna Drosophila Resource Center, InDroso and the Developmental Studies Hybridoma Bank for providing fly stocks and antibodies. We also thank the Microscopy Rennes Imaging Center-BIOSIT facility. We thank S.Luschnig, M. Hollmann and members of R.L.B.’s laboratory for critical reading of the manuscript. This work was supported in part by the ARED program from the Région Bretagne, La Ligue contre le Cancer-Equipe Labellisée (R.L.B.) and the Association Nationale de la Recherche et de la Technologie programme PRC Vie, santé et bien-être CytoSIGN (ANR-16-CE13-004-01 to R.L.B.).

## Author contributions

Conceptualization, T.E.d.B and R.L.B.; Methodology, T.E.d.B and R.L.B.; Investigation, T.E.d.B and R.L.B.; Formal Analysis, T.E.d.B.; Visualization, T.E.d.B and R.L.B.; Writing – Original Draft, T.E.d.B and R.L.B.; Writing – Review & Editing, T.E.d.B and R.L.B.; Funding Acquisition, R.L.B.; Supervision, R.L.B.

## Declaration of interest

The authors declare no competing interests

## KEY RESOURCES TABLE

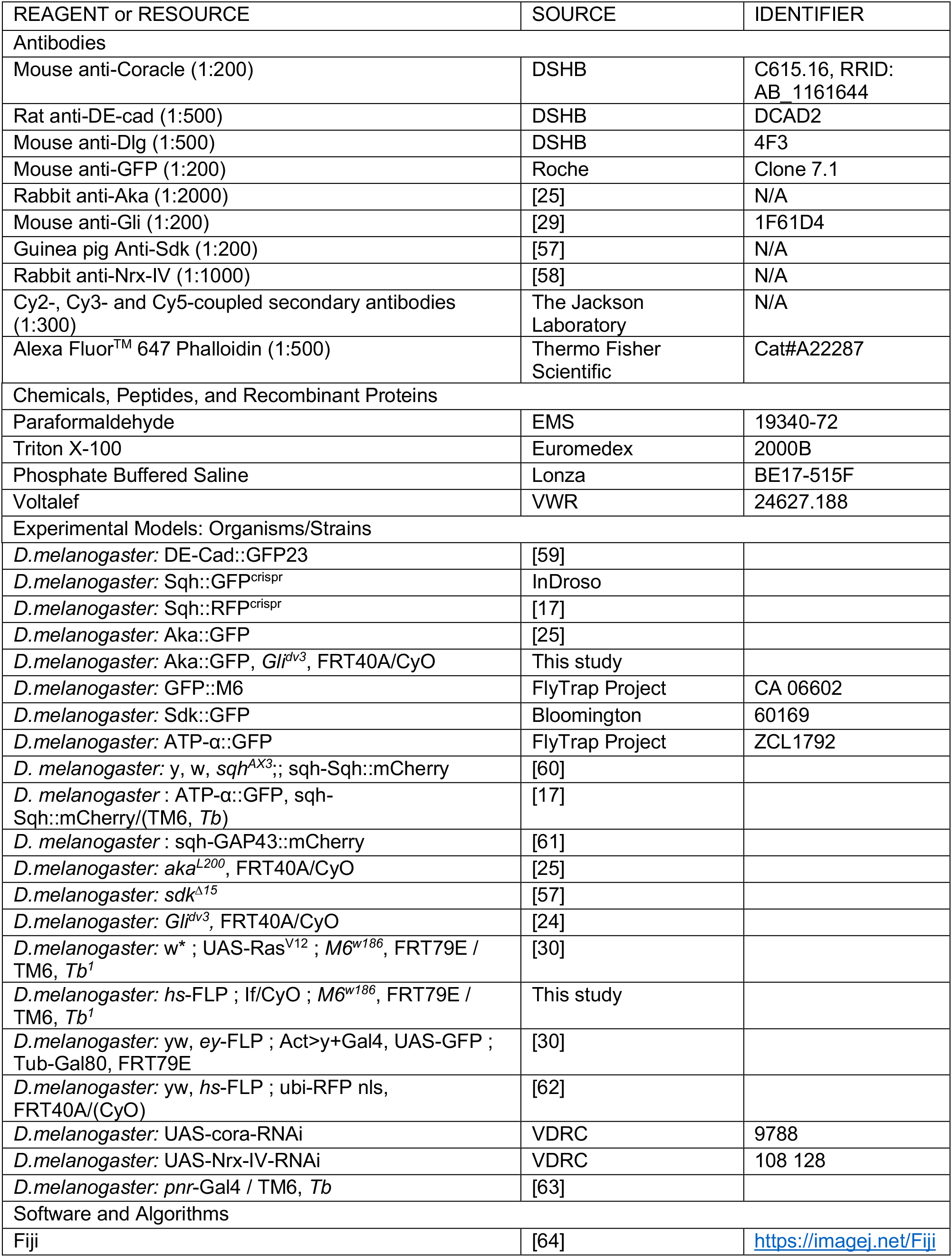
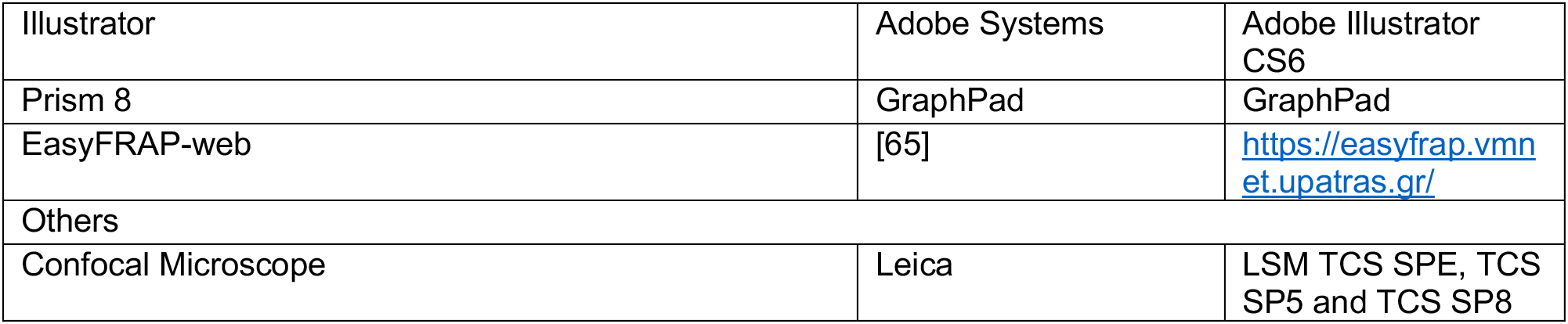

## Drosophila genotypes

**Figure 1**

(B) sqh∷RFP^crispr^ / Sdk∷GFP

(D) sqh∷RFP^crispr^ ; ; M6∷GFP

(E) sqh∷RFP^crispr^ ; Aka∷GFP

**Figure 2**

(A-D) *hs*-FLP ; *aka*^*L200*^, FRT40A / ubi-RFP nls, FRT40A ; ATP-α∷GFP, sqh-Sqh∷mCherry / +

**Figure 3**

(A) *w*^*118*^

(B) *sdk*^*Δ15*^

(C-E) *hs*-FLP ; *aka*^*L200*^, FRT40A / ubi-RFP nls, FRT40A

(F) *hs*-FLP ; Act>y+Gal4, UAS-GFP ; *M6*^*w186*^, FRT79E / Tub-Gal80, FRT79E

**Figure 4**

(A) *hs*-FLP ; *Gli*^*dv3*^, FRT40A/ ubi-RFP nls, FRT40A

(B-C) *hs*-FLP ; Aka∷GFP, *Gli*^*dv3*^, FRT40A/ ubi-RFP nls, FRT40A

**Figure 5**

(A, C, D, E, G and I) *hs*-FLP ; *aka*^*L200*^, FRT40A / ubi-RFP nls, FRT40A

(B) *hs*-FLP ; *aka*^*L200*^, FRT40A / ubi-RFP nls, FRT40A ; ATP-α∷GFP, sqh-Sqh∷mCherry/ +

**Figure 6**

(A-E) *hs*-FLP ; *aka*^*L200*^, FRT40A / ubi-RFP nls, FRT40A ; ATP-α∷GFP, sqh-Sqh∷mCherry / +

**Figure 7**

(A-C) ; ; UAS-cora-RNAi / *pnr*-Gal4

## Method details

### Immunofluorescence

Pupae aged for 16h30 to 19h after puparium formation (APF) were dissected using Cannas microscissors in 1X Phosphate-Buffered Saline (1X PBS, pH 7.4) and fixed 15 min in 4% paraformaldehyde at room temperature [66]. Following fixation, dissected nota were permeabilized using 0.1% Triton X-100 in 1X PBS (PBT), incubated with primary antibodies diluted in PBT for 2 hours at room temperature. After 3 washes of 5 minutes in PBT, nota were incubated with secondary antibodies diluted in PBT for 1 hour, followed by 2 washes in PBT, and one wash in PBS, prior mounting in 0,5% N-propylgallate dissolved in 90% glycerol/PBS 1X final.

### Live-imaging and image analyses

Live imaging was performed on pupae aged for 16h30 APF at 25°C. Pupae were sticked on a glass slide with a double-sided tape, and the brown pupal case was removed over the head and dorsal thorax using microdissecting forceps. Pillars made of 4 and 5 glass coverslips were positioned at the anterior and posterior side of the pupae, respectively. A glass coverslip covered with a thin film of Voltalef 10S oil is then placed on top of the pillars such that a meniscus is formed between the dorsal thorax of the pupae and the glass coverslip [67]. Images were acquired with a LSM Leica SPE, SP5 or SP8 equipped with a 63X N.A. 1.4. and controlled by LAS AF software. Confocal sections (z) were taken every 0.5 μm. For figures representation, images were processed with Gaussian Blur σ = 1.1. All images were processed and assembled using Fiji software [64] and Adobe Illustrator.

### Fluorescence recovery after photobleaching

FRAP experiments were performed in pupae expressing Aka∷GFP, Venus∷Aka together with RFP-nls used as a marker of *Gli*^*dv3*^ cells. Aka∷GFP and Venus∷Aka were bleached (488 nm laser at 60% power, 2 iterations of 1.293 s, square ROI of 2μm*1.5μm) using a LSM Leica SP5 or SP8 equipped with a 63X N.A. 1.4 PlanApo objective. Confocal stacks were acquired every 30s or 2 min before and after photobleaching on 13 z steps to compensate for movement in z during acquisition. FRAP experiments were performed in pupae expressing ATP-α∷GFP with sqh∷RFP^crispr^ to localize AJ level. ATP-α∷GFP was bleached (488 nm laser at 60% power, 2 iterations of 1.293 s, 2.5μm*1.5μm) using a LSM Leica SP5 or SP8 equipped with a 63X N.A. 1.4 PlanApo objective. Confocal stacks were acquired every 30s or 2 min before and after photobleaching on 6 or 14 z steps to compensate for movement in z during acquisition.

### Nanoablation

Laser ablation was performed on live pupae aged for 16h to 19h APF using a Leica SP5 confocal microscope equipped with a 63X N.A. 1.4 PlanApo objective. Ablation was carried out on epithelial cell membranes at SJ level with a two-photon laser-type Mai-Tai HP from Spectra Physics set to 800 nm and a laser power of 2.9W. LAS AF parameters are laser trans 50%, gain 70%, offset 35% and 1 iteration of 1.293 s.

## Quantification and statistical analysis

### Signal recovery upon photobleaching

In all FRAP experiments, ROI1 corresponds to the photobleached area, ROI2 to the area used as a control for general photobleaching of the sample across time and ROI3 to the background. ROI1 and RIO3 had the same dimensions in all FRAP experiments (2μm*1.5μm for Aka∷GFP and Venus∷Aka ; 2.5μm*1.5μm for ATP-α∷GFP).

For Aka∷GFP and Venus∷Aka, a sum slice of 8 z (4μm in total) was applied in order to collect the entire signal. For Aka∷GFP, ROI2 was considered as the fluorescence of the entire field. For Venus∷Aka, 3 unphotobleached vertices were measured and the mean was taken as the value for ROI2.

For ATP-α∷GFP, a sum slice, of 2 z (1μm in total) of the plan just below AJ marked by sqh∷RFP^crispr^ and the one below, was applied to avoid z movements. ROI2 was considered as the fluorescence of the entire field.

Then, we used EasyFRAP-web [65] for all data processing and to extract normalized value. The I(t) which correspond to the fluorescence intensity across time was calculated as described:

First, the fluorescence intensity is corrected by subtracting the background:

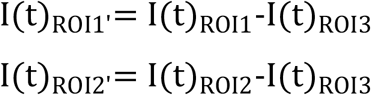

Then, values of fluorescence intensity are normalized in this way:

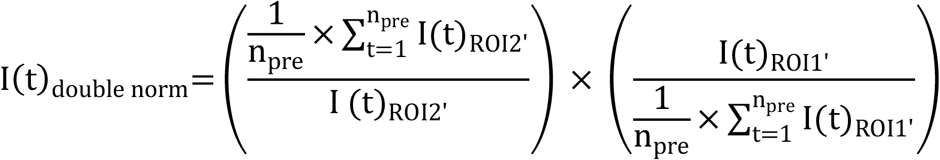

We decided to take value from full scale normalization in order to have a curve starting from 0:

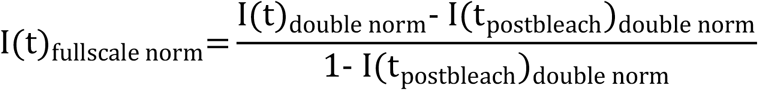

Recovery curves were then fitted assuming a one-phase exponential association equation using the GraphPad Prism 8.0 software:

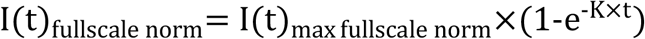

We admitted that I(t)_max fullscale norm_ is the maximum value based on the fitted curve for the post-bleach time (20 min for Aka∷GFP, Venus∷Aka and 18 min for ATP-α∷GFP) and it is considered as the immobile fraction. t_1/2_ was deduced when I(t)_fullscale norm_=0,5×I(t)_max fullscale norm_ which occured at 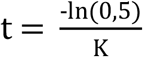.

### Fluorescence vertex analysis

Sum slices were applied to different experiments. A line of 10 pixels width was used to measure Anakonda signal in WT and *Gli* ^*dv3*^ mutant vertex. Using the same width, line were draw to extract background fluorescence signal and the background signal was subtracted to each vertex quantification. After, data were normalized between 0 and 1 to allow visual representation with 1 corresponding to the highest signal of Anakonda at vertex in each experiments analyzed and 0 the lowest.

### Line scan fluorescence analysis

Maximal projection were applied and a 20 pixels width line was drawn from apical part to basal part (Figure 5F-H’) or from vertex 1 to vertex 2, spanning the BCJ (Figures 5I’, 7B’ and C’). Then gray value was plotted across length of the line.

### Statistical tests

All information concerning the statistical details are provided in the main text and in figure legends, including the number of samples analyzed for each experiment. Scattered plots use the following standards: thick line indicate the means and errors bars represent the standard deviations. Boxplots with connected line use the following standards: dots represent mean and the total colored areas show SD.

Statistical analyses were performed using the GraphPad Prism 8.0 software. The Shapiro-Wilk normality test was used to confirm the normality of the data and the F-test to verify the equality of SD. The statistical difference of Gaussian data sets was analyzed using the Student unpaired two-tailed t test. For contingency analysis (Figures 2E and S2C), a Fisher’s exact test was used. Statistical significances were represented as follow: p value > 0.05 NS (not significant) and p value ≤ 0.0001 ****.

## Notes

### Competing Interest Statement

The authors have declared no competing interest.

