## Supplemental Information for "Interplay between Anakonda, Gliotactin and M6 for tricellular junction assembly and anchoring of septate junctions in *Drosophila* epithelium"

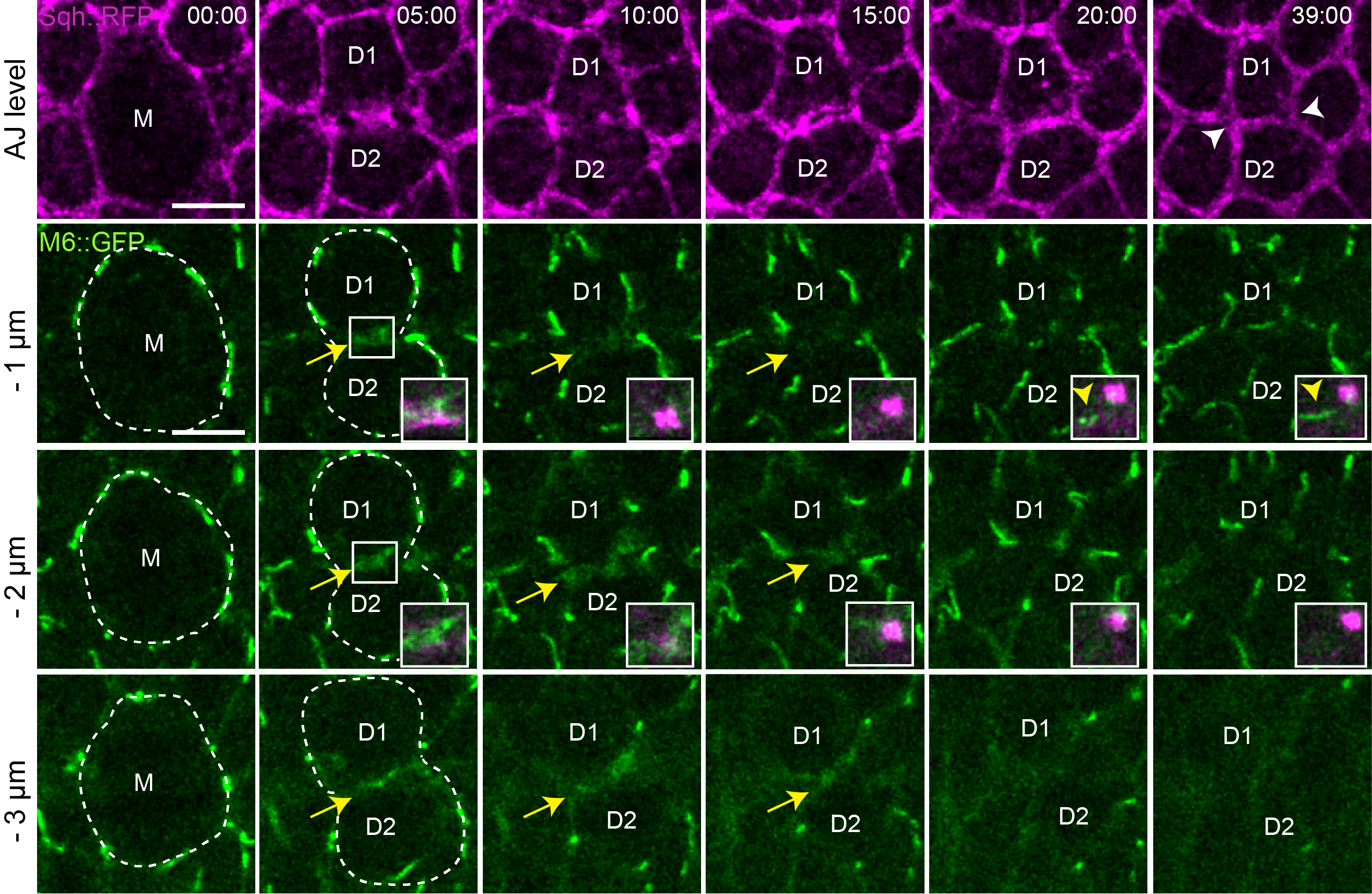

**Figure S1:** **Spatiotemporal analysis of M6::GFP assembly during epithelial cytokinesis**

Time-lapse imaging of Sqh::RFP^crispr^ (magenta, AJ) and M6::GFP (tSJ). The mother cell is represented by the M and the daughters by D1 and D2. The white dashed line highlights the divided cell and the two new cells. Yellow arrows show M6::GFP foggy signal at the future daughter-daughter cell interface. Yellow arrowheads show the signal of M6::GFP appearance close to the midbody. White arrowheads show the two new TCJs formed. White squares show high magnification of M6::GFP close to the midbody (magenta). Time is min:sec with t = 0 corresponding to the anaphase onset. Distances correspond to the position relative to the plane of AJ labeled with Sqh::RFP. The scale bars represent 5µm.

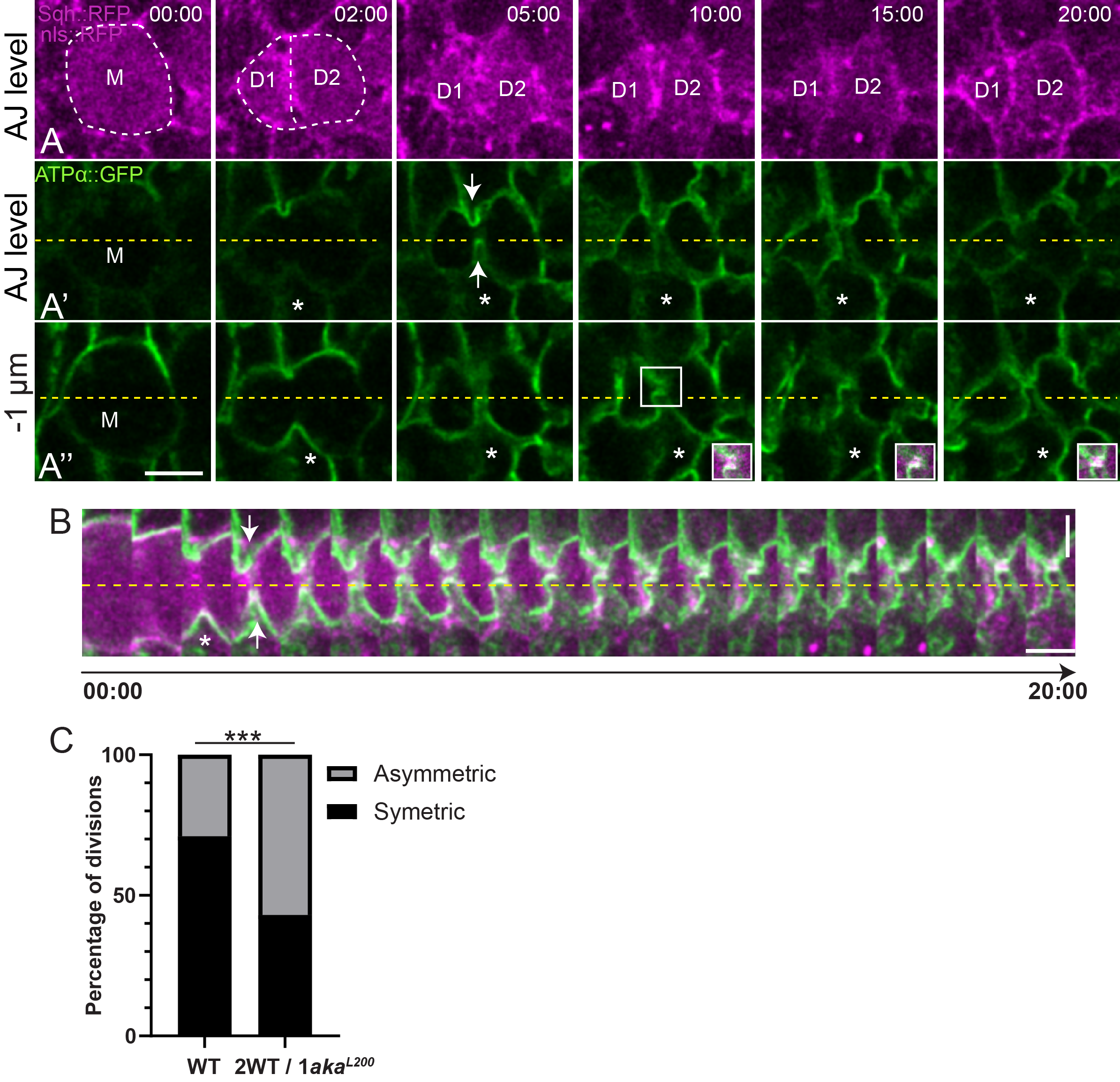
**Figure S2:** **Loss of integrity of tSJ from a neighboring cell impaired FLP formation during cell epithelial cytokinesis**

(A-B) Time-lapse imaging of FLP using Sqh::RFP^crispr^ (magenta, AJ) and ATPα::GFP (green, SJ) in wild type dividing cell between a wild type (marked by nls::RFP) and an *aka^L200^* cell (loss of nls::RFP), from plane view (A-A’’) and with a kymograph representation (B). The mother cell is represented by the M and the daughters by D1 and D2. The white dashed lines highlight the divided cell and the two new cells. White arrows indicate FLP formation at SJ level. White asterisk shows the *aka^L200^* cell. Yellow dashed lines define symmetry axis of the cell. White squares show high magnification of midbody (magenta) connected to FLPs. (C) Histogram representing the number of symmetric (Black) and asymmetric (Grey) FLP formation during cytokinesis in wild type (symmetric = 71% ; asymmetric = 29% ; n= 14 divisions, 3 pupae) and in wild type with a finger coming from a wild type cell and a finger coming from an *aka^L200^* cell (symmetric = 43% ; asymmetric = 57% ; n= 14 divisions, 3 pupae). *** p < 0.001, Fisher’s exact test. The horizontal scale bar represents 5µm (A), the vertical scale bar represents 5µm (B) and the horizontal scale bars represent 1min (B).

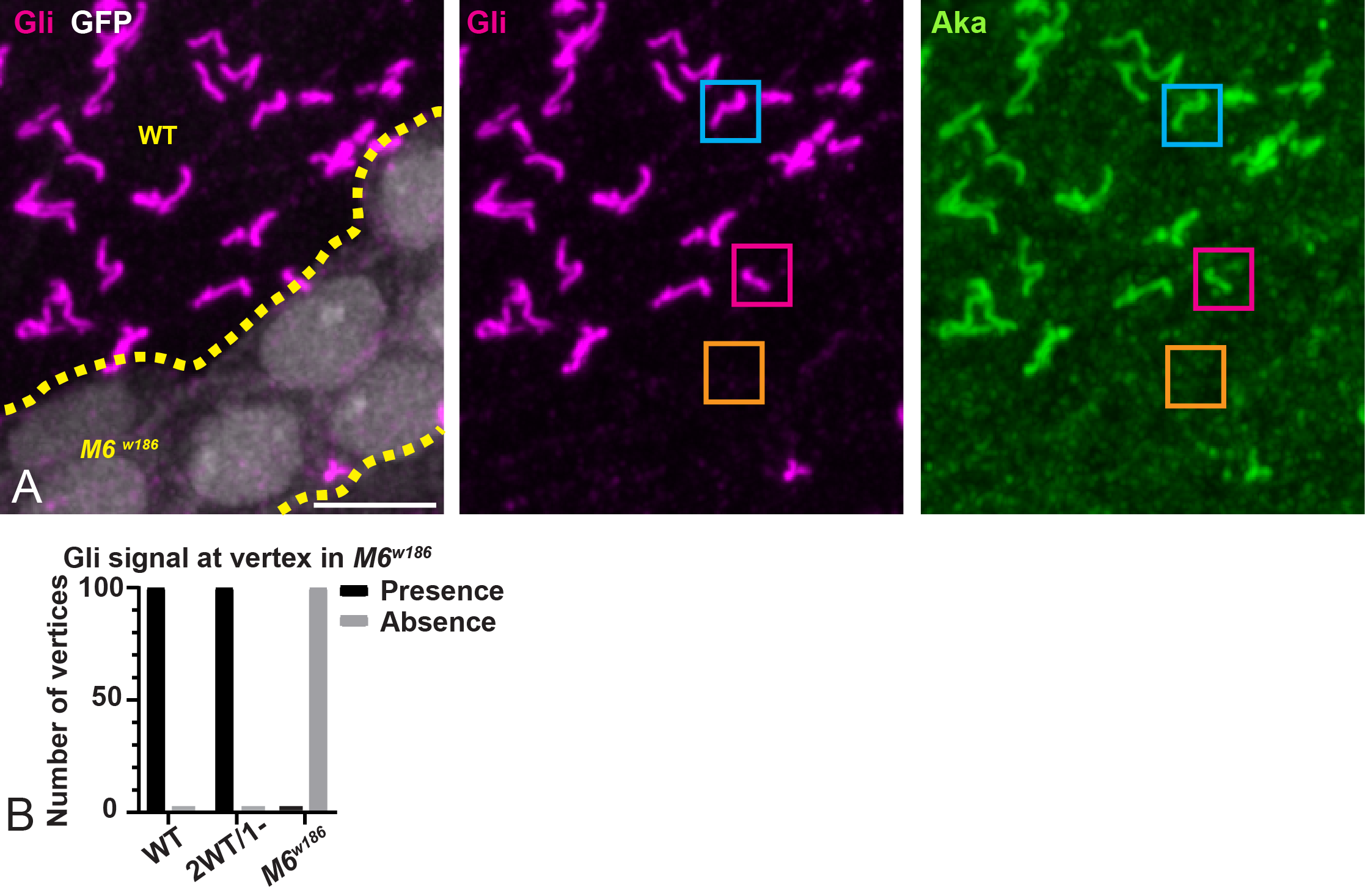
**Figure S3: M6 is required to localize Gli at the tSJ**

(A) Notum at 16h30 APF stained for Gli (anti-Gli, magenta), Aka (anti-Aka, green) after heat-shock to induce clone of wild type (GFP negative) and *M6* cells (GFP positive). Gli is enriched at the TCJ at wild-type vertex (blue square) or a vertex between 2 wild type and 1 *M6* cells but disappeared at a vertex of three *M6^w186^* cells (orange square). (B) Histogram representing the percentage of presence (black ) or absence (gray) of Gli at the vertex between 3 wild type cells (presence n= 100, absence n = 0), 2 wild type and 1 *M6^w186^* cells (presence n= 100, absence n = 0) or 3 *M6^w186^* cells (presence n= 0, absence n= 100) (n= 100 vertices, > 5 pupae). The scale bar represents 5µm. Images are maximum projection.

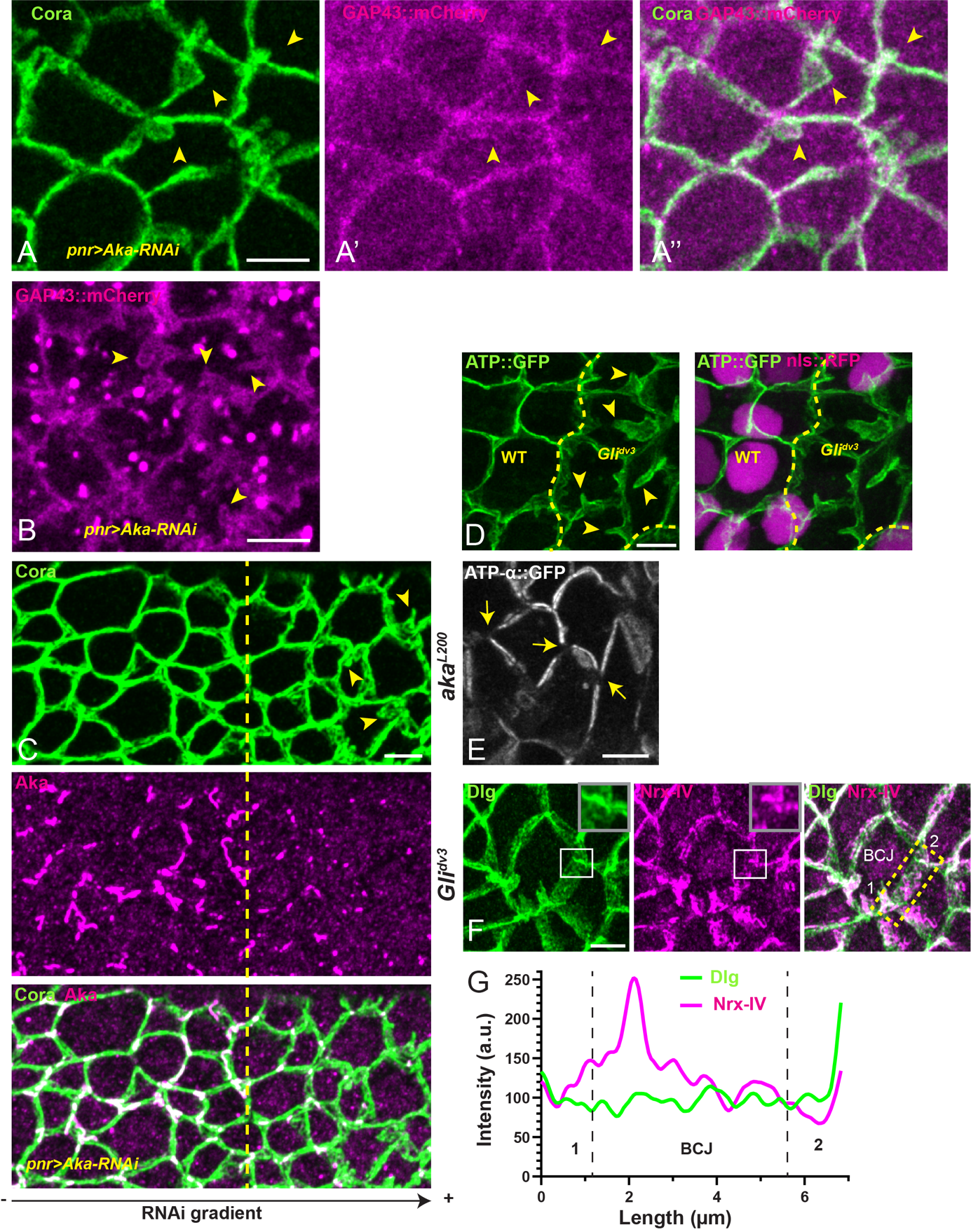

**Figure S4:** **tSJ ensures SJ shape integrity and SJ core components anchoring at the vertex**

(A-D) SJ and membrane morphology analysis via core components Cora (anti-Cora; A and C), GAP43::mCherry (A-B), Aka (anti-Aka; C) and ATP-α::GFP (D) in wild type and upon knockdown of Aka by RNAi approach (A-C) or *Gli^dv3^* cells (D). (A) Showed GAP43::mCherry tagged membrane after fixation whereas (B) shows live imaging of GAP43::mCherry tagged membrane in Aka RNAi area. Yellow arrowheads indicate micrometric size SJ/membrane deformations. The dashed yellow line separates wild type from Aka knockdown cells. (E) Localization of ATP-α::GFP in *Gli^dv3^* cells. Yellow arrows show loss of ATP-α::GFP at vertices. (F) Localization of Dlg (anti-Dlg, green) and Nrx-IV (anti-Nrx-IV, magenta) in *Gli^dv3^* cells after maximal projection. White squares show high magnification of a vertex in *Gli^dv3^* cells. Yellow dashed rectangle shows the line scan used between two vertices represented by 1 and 2 to obtain (G) Plot representing Dlg (green line) and Nrx-IV (magenta line) signals as a function of the length of cell-cell boundary.

The scale bars represent 5µm.

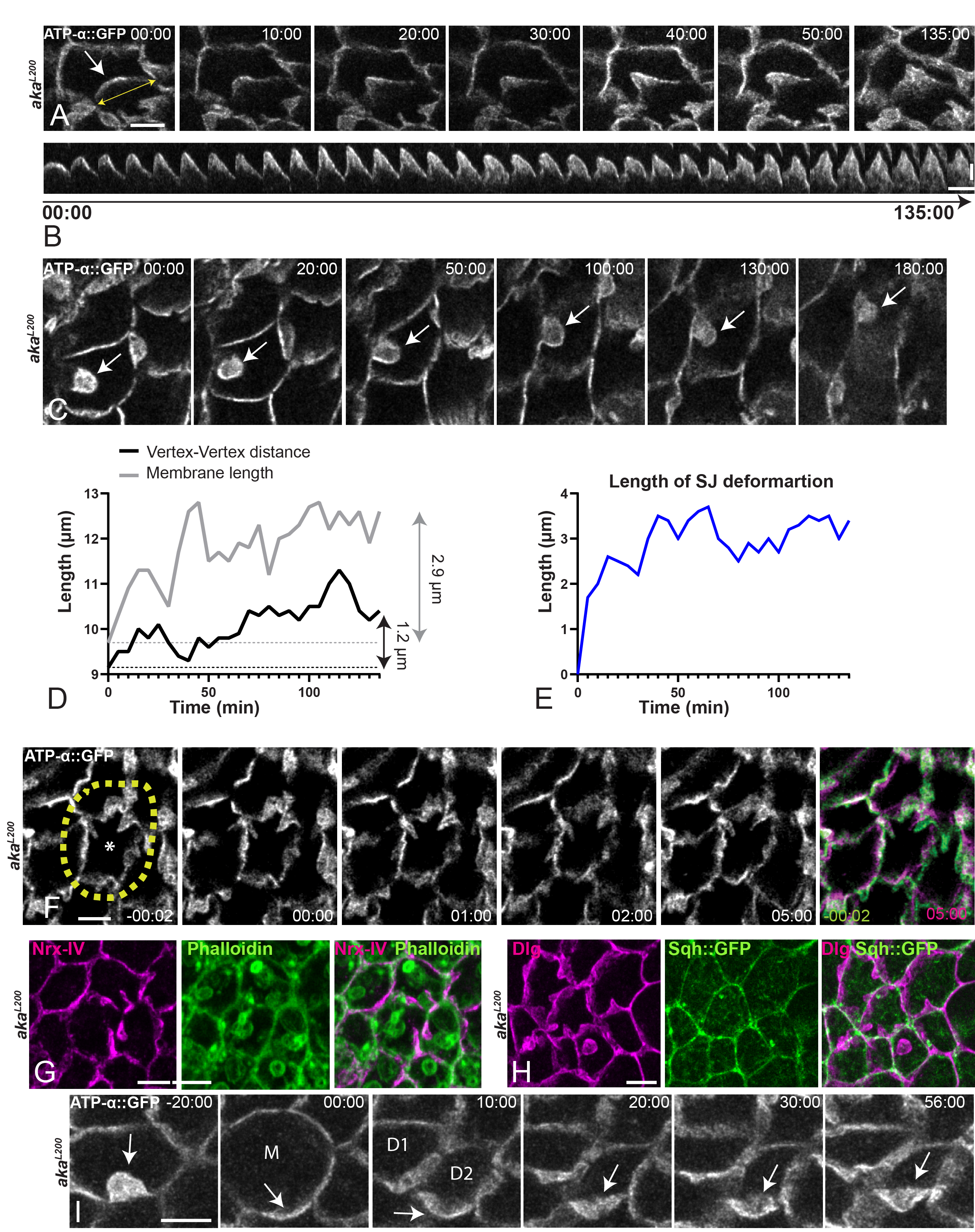

**Figure S5:** **SJ deformations appear slowly, are not under mechanical tension or enriched in actomyosin cytoskeleton and are deformable during cytokinesis**

(A-B) Time-lapse imaging of ATP-α::GFP positive SJ deformation appearance in *aka^L200^* cells from planar view (A) and with a kymograph representation (B). (C) ATP-α::GFP positive SJ deformation remains at the same Z level during at least 180min (D-E). Plot of the vertex-vertex distance (black line) or total ATP-α::GFP marked membrane (gray line) in µm (C) or ATP-α::GFP positive SJ deformation length (blue line) in µm (D) across time (min) of the experiment depicted in (A-B). (F) Bi-photonic laser-based nano-ablation all around one *aka^L200^* cell expressing ATP-α::GFP (white). Green and magenta pictures represent SJ deformations 2s before and 5mins after ablation respectively. Yellow dashed line shows the ablation area. (G-H) Notum between 16h and 18h APF stained for Nrx-IV (anti-Nrx-IV, magenta; F), Dlg (anti-Dlg, magenta; G), Phalloidin (green; F) or expressing Sqh::GFP (green; G) in *aka^L200^* cells. (I) Time-lapse imaging of ATP-α::GFP positive SJ deformation in *aka^L200^* cells. The mother cell is represented by the M and the daughters by D1 and D2. White arrows show the ATP-α::GFP positive SJ deformations. Time is min:sec with t= 0 corresponding to the SJ deformation onset (A) or anaphase onset (I). The horizontal scale bars represent 5µm (A, C, F, G, H and I) or 4 mins (B). The vertical scale bar is 2 µm (B).

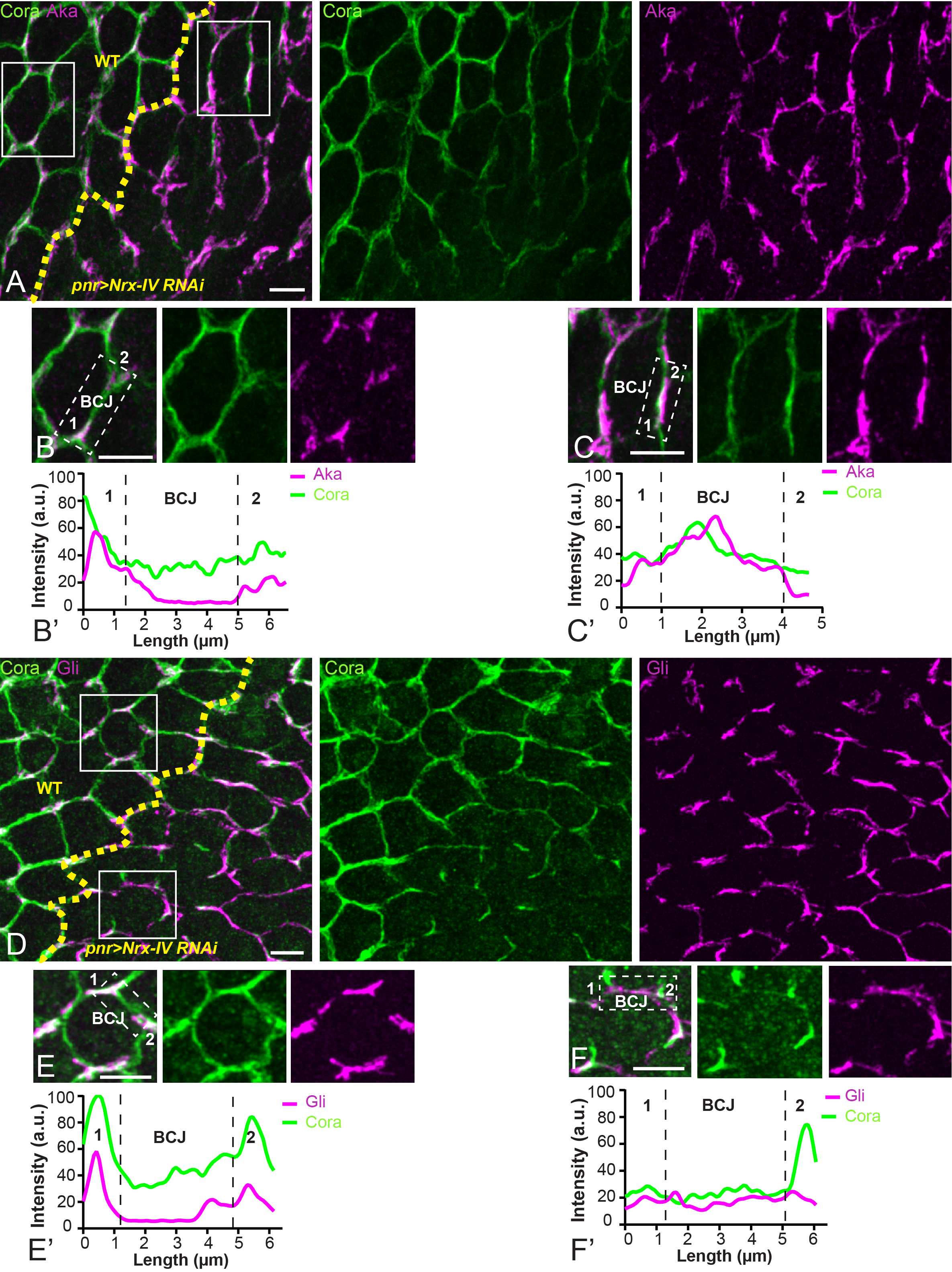

**Figure S6: The activity of SJ core component is required to restrict Aka and Gli localization at vertex**

(A-C, D-F) Localization of Aka (anti-Aka, magenta; A-C) or Gli (anti-Gli, magenta; D-F) in cells marked by Cora (anti-Cora, green) in wild type and cells expressing UAS::Nrx-IV RNAi under *pnr*::Gal4 control. The dashed yellow line separates wild type and Nrx-IV-RNAi cells. Aka and Gli spread at the BCJ upon knockdown of Nrx-IV. White squares (A and D) show magnification (B-C and E-F) of wild type and knockdown cells for Nrx-IV. White dashed rectangles show the line scan used to obtain (B’-C’ and E’-F’) between two vertices represented by number 1 and 2. (B’-C’ and E’-F’) Plots representing Cora (green line) and Aka/Gli (magenta line) signals as a function of the length of cell-cell boundary in wild type and Nrx-IV-RNAi cells respectively. The scale bars represent 5µm.
